# Fluorogenic EXO-Probe Aptamers for Imaging and Tracking Exosomal RNAs

**DOI:** 10.1101/2021.08.18.456703

**Authors:** Emily E. Bonacquisti, Scott W. Ferguson, Natalie E. Jasiewicz, Jinli Wang, Adam D. Brown, Daniel P. Keeley, Michelle S. Itano, Juliane Nguyen

## Abstract

Small extracellular vesicles (sEVs), or exosomes, play important roles in physiological and pathological cellular communication. sEVs contain both short and long non-coding RNAs that regulate gene expression and epigenetic processes. Studying the intricacies of sEV function and RNA-based communication requires tools capable of labeling sEV RNA. Here we developed a novel genetically encodable reporter system for tracking sEV RNAs comprising an sEV-loading RNA sequence, termed the EXO-Code, fused to a fluorogenic RNA Mango aptamer for RNA imaging. This fusion construct allowed the visualization and tracking of RNA puncta and colocalization with markers of multivesicular bodies; imaging RNA puncta within sEVs; and quantification of sEVs. This technology represents a useful and versatile tool to interrogate the role of sEVs in cellular communication via RNA trafficking to sEVs, cellular sorting decisions, and sEV RNA cargo transfer to recipient cells.

## Introduction

Small extracellular vesicles (sEVs), also known as exosomes, play important roles in physiological and pathological cellular communication^1–5^. sEVs contain numerous short and long non-coding RNAs that regulate multiple aspects of gene expression, including the epigenetic processes that modulate cell fate, polarization, and morphogenesis^3, 6–11^. Despite the important roles played by exosomal RNAs, relatively little is known about mechanisms of exosome-RNA packaging, trafficking, and fate upon uptake by target cells. This is primarily because there are no tools to directly label sEV RNA cargoes in cells. While fluorogenic RNA aptamers that dramatically increase the fluorescence of small molecule fluorophores upon binding are described, they do not enrich into sEVs and localize non-specifically in cells^12, 13^. Other methods to track sEV RNAs require the exogenous introduction of RNAs chemically pre-labeled with fluorophores. However, these do not allow the tracking of RNA sorting into sEVs from their early biogenesis to secretion, and therefore are not genetically encodable.

Given the central importance of exosomal RNAs in dictating cellular behavior, there is therefore a need for sensitive and specific exosomal RNA imaging tools to determine how cells use sEVs and their RNA cargoes in communication. For example, in aggressive cancers, sEVs travel to distant organs and prime distant sites for metastasis using their RNA cargo; however the exact mechanism by which they do this remains unknown^14–17^. It is difficult to label RNAs while maintaining their biological function, and access to sEV RNAs is further complicated by sEVs harboring very few copies of a given RNA species, oftentimes less than one miRNA copy per vesicle^18^. Elucidating the trafficking and processing of exosomal RNA demands both novel markers capable of tracking the intercellular movement of exosomal RNA and enhanced loading of RNAs into exosomes.

Most existing sEV labeling methods modify the exosomal surface through lipid labeling or membrane fusion proteins. Tracking sEVs with lipid-based fluorophores is hampered by non-specific lipid-lipid transfer to proximal cell membranes, potentially confounding readouts^19–21^. Other significant advances related to sEV labeling rely on the generation of fusion proteins, which have limitations. For example, luciferase-tagged EVs can be used to monitor EV kinetics and dynamics *in vivo* but the surface modification can have varying effects on the native biodistribution^22, 23^.

A pH-sensitive GFP derivative, pHlourin, was recently engineered as a tetraspanin tag, allowing the monitoring of endosomal fusion events of pHlourin-positive vesicles^24–26^. Later iterations of this tool yielded a pH-sensitive mScarlet fluorophore capable of fluorescing under different pH conditions^25^. While these and other biochemical tools used to label sEVs have advanced the field, they are limited to tracking the sEV lipid membrane and not their contents. Also, addition of surface proteins to sEV membranes may alter downstream biological targeting and uptake mechanisms^27, 28^. A novel system is needed to track the movement of not only sEVs but also their RNA cargoes.

Here we developed a genetically encodable reporter system for tracking sEV RNAs. This novel encodable reporter system comprises a fluorogenic RNA Mango aptamer fused to a newly identified RNA sequence, termed an EXO-Code, that substantially enriches into sEVs. The EXO-Code showed a greater number of accelerated linear trajectories towards the cell membrane compared to a random RNA sequence and bound to a distinct set of cellular proteins. Moreover, we identified the structural characteristics of an EXO-Code/Mango fusion construct crucial for sEV enrichment while maintaining binding to thiazole orange (TO) for imaging and tracking of sEV RNAs. Using this construct, we visualized bright RNA puncta and their trafficking within cells, imaged RNA puncta within sEVs, and quantified sEVs.

## Results

### Identifying biomimetic zip code-like RNA sequences that mediate sorting to sEVs

We developed a novel approach to screen for biomimetic RNA “zip codes”, which we term EXO-Codes, that are highly enriched into sEVs (**Fig. 1a**). Conventional approaches screen for sequences against isolated protein targets outside their native cellular environment^29^. However, the sorting of nucleic acids into sEVs is a highly complex and poorly understood process where the identity of potential sorting proteins remains unclear^6, 16, 30, 31^. To address this, we subjected an RNA library to the native intracellular environment of living cells to harness the endogenous cellular trafficking machinery for the active sorting of EXO-Codes to sEVs. By maintaining the native intracellular environment, cellular sorting mechanisms were preserved and targeted. By using chemically synthesized RNA sequences of high diversity, we introduced sequences with the potential to outperform natural miRNA and mRNA in their ability to sort to sEVs. Thus, this method selected for sequences that show high sEV enrichment compared to existing approaches that rely on exploiting endogenous miRNA motifs within extracellular miRNAs^32, 33^.

**Figure 1.**
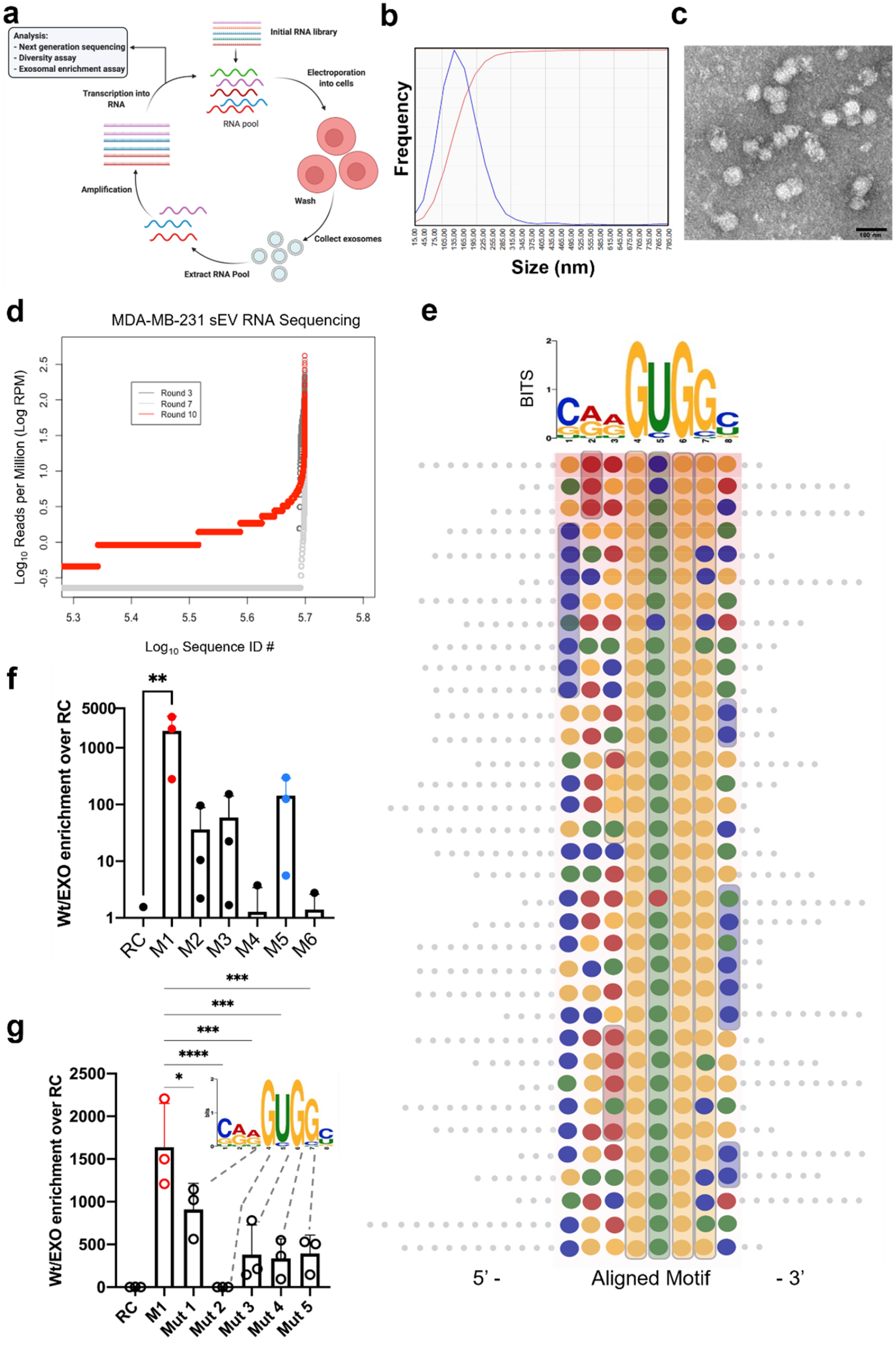
Bioinformatic and *in vitro* characterization of RNA EXO-Codes in MDA-MB-231 cells. (**a**) Experimental workflow of the EXO-Code selection procedure. Each complete cycle through the procedure is represented as one round, and NGS sequencing and analysis was performed at rounds 3, 5, 7, and 10. Each collected portion of extracted sEV RNA was amplified and re-introduced into cells to use the endogenous cellular sorting machinery to uncover the top EXO-Codes**. (b)** Representative MDA-MB-231 sEV NTA particle size distribution. **(c)** Representative transmission electron microscopy images of MDA-MB-231 sEVs, scale bar set at 100 nm. **(d)** Visualization of next-generation sequencing data compiled as sequence counts versus log-normalized reads per million, with each dot representing an individual and unique sequence. Round 3 plotted in black, round 7 in gray, and round 10 plotted in red. The overlap in sequence ID number at the upper right of the graph indicates the same unique sequences enriching into sEVs in higher amounts as the selection procedure moved to later rounds. **(e)** Schematic depicting MEME-predicted motif sites in the top 50 sequences, color-coded according to IACUC standards. Eight nucleotide RNA motif required for EXO-Code function (top of panel e). The red shading indicates the NGS rank of sequences with the highest RPM values. This visualization was used to choose the top sequences tested in panel f. **(f)** Quantitative RT-PCR endpoint enrichment analysis of random control and EXO-Codes M1-M6, data presented as mean ± standard deviation. **i.** Quantitative RT-PCR endpoint enrichment of mutated M1 nucleotides, data presented as mean ± standard deviation. Experiments f and g were completed in triplicate with representative experiment shown. hMSC data available in Supplementary Fig. 1. Statistical comparisons were performed using one-way ANOVA \*\*\*\**p* < 0.0001, \*\*\**p* < 0.001, \*\**p* < 0.01, \**p* < 0.05.

We used sEVs from the MDA-MB-231 triple-negative breast cancer cell line as a model for EXO-Code screening. The *de novo* EXO-Code discovery method was also applied to a second cell type, human bone-marrow-derived mesenchymal stem cells (hMSCs), to demonstrate generalizability of the method for obtaining cell-type specific EXO-Codes (**Supplementary Fig. 1**). To select for RNA sequences enriched into sEVs, MDA-MB-231 cells were electroporated with a 10^12^-diverse RNA library, followed by several washes to remove extracellular RNA. To allow EXO-Codes to sort to sEVs, cells were grown in exosome-depleted medium before collecting secreted sEVs by differential ultracentrifugation using established protocols^34^. MDA-MB-231 and hMSC sEVs had a mean diameter of ∼105 nm by nanoparticle tracking analysis (NTA) and ∼100 nm by transmission electron microscopy (TEM) (**Fig. 1b,c and Supplementary Fig. 1b,c**).

Enriched RNA EXO-Code reads were analyzed by next-generation sequencing (NGS; Illumina MiSeq2000). At later rounds of the sEV RNA sequencing procedure the overall sequence diversity drastically decreased, corresponding to a sharp increase in average reads per million (RPM) of the top sequences at the final round of selection, confirming selection success (**Fig. 1d, Supplementary Fig. 1a**).

The top 25 sequences (**Fig. 1e, f**) were ranked to resolve the top five candidates displaying the highest sEV enrichment. After validation with endpoint quantitative (q)PCR, the top performing code, henceforth referred to as “M1” (MDA-MB-231 rank order 1), was approximately 3000-fold more enriched in sEVs than a random control of the same nucleotide content (**Fig. 1f)**. Furthermore, the M1 sequence enriched >1000-fold over a previously reported motif thought to preferentially enrich into exosomes (**Supplementary Fig. 2**). The hMSC top performer designated as “S1 (stem cell rank order 1), and was enriched ∼2000-fold more into sEVs compared with the random control sequence (**Supplementary Fig. 1e)**. As a next step, we sought out to determine a conserved motif in the most enriched sequences and discovered an 8-nucleotide RNA motif, termed the EXO-Code, that was present in 98.5% of the top 200 sequences (**Fig. 1e, Supplementary Fig. 1d)**.

To assess if the EXO-Code enriched into sEVs in a sequence-specific manner, mutations were introduced at key positions in the EXO-Code motif, i.e., positions that displayed >85% of the position-dependent letter probability, here GUGG and the adjacent C, were considered key positions. As such, we mutated those five nucleotide residues individually and reassessed for sEVs enrichment in MDA-MB-231 cells and hMSCs. Mutation of the first G >> U decreased sorting by 55.6%, the U>>G mutation decreased sorting efficiency by 98.07%, while mutations at the last three positions decreased sorting by 77.4% in MDA-MB-231 cells (**Fig. 1g)**. For hMSCs, all single nucleotide mutations within 5 key positions in the S1 sequence abolished >99% of EXO-Code enrichment (**Supplementary Fig. 1f**). We concluded that EXO-Code sorting is likely to be highly sequence specific (**Fig. 1g, Supplementary Fig. 1f**). We next sought to understand the interplay between intracellular sorting proteins and subcellular locomotion of our novel EXO-Codes using MDA-MB-231 cells as our model cell line.

### EXO-Codes use active transport pathways to move toward the plasma membrane

Some specific miRNAs are sorted over other types of miRNAs into sEVs, but the exact mechanisms of this specificity are not fully understood^7, 16, 32, 33^. We hypothesized that the EXO-Codes were also differentially trafficked to the cellular membrane compared to a non-specific RNA control sequence for packaging into sEVs. We then tested this hypothesis by tracking the movement of Alexa-488 tagged EXO-Codes and a non-specific RNA control in MDA-MB-231 cells in real time using high-resolution live cell imaging confocal microscopy.

Control RNA was visibly sequestered within intracellular pockets, with >80% being immobile, whereas our EXO-Codes moved quickly and in a directed fashion within the cytosol (**Fig. 2a,b; Supplementary Movies 1 and 2**). By quantifying the kinetics of the RNA puncta, we established that the EXO-Code RNA traveled approximately two-fold further and in a more linear fashion than the control RNA sequence (**Fig. 2c,d)**. EXO-Code puncta were also located close to the plasma membrane in a less sequestered fashion than the control (**Fig. 2e,f)**. 93.3% of EXO-code puncta were mobile compared to only 16.9% of the control puncta (**Fig. 2e)**. In combination with the increased proximity to the plasma membrane **(Fig. 2f)**, these data could indicate that the EXO-Code puncta were more likely engaged with mechanisms enabling them to being exported out of the cell.

**Figure 2.**
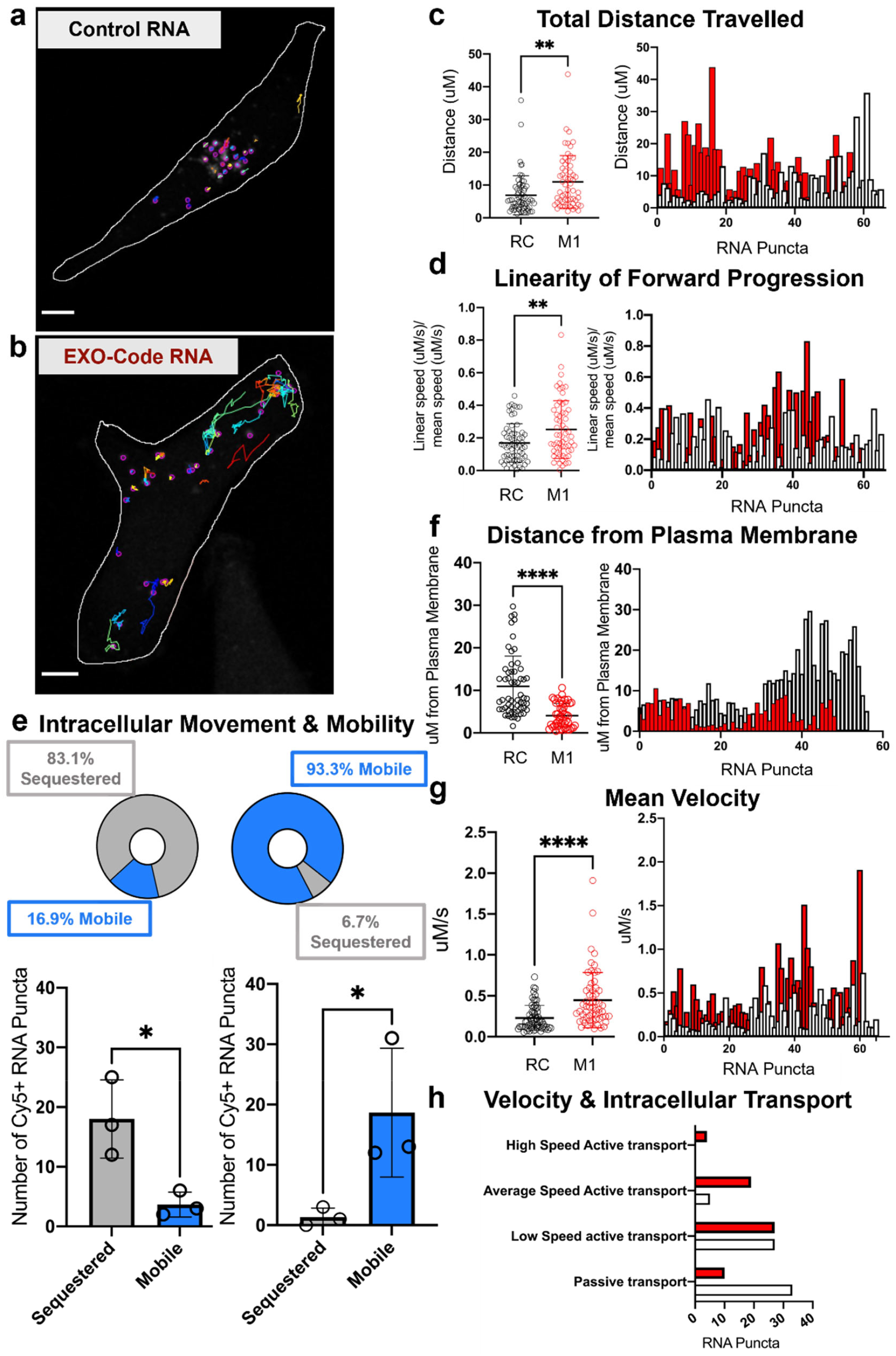
Live-cell imaging of EXO-Code movement in MDA-MB-231 cells. **(a-b)** Representative video stills of AF488-RNA puncta annotated with Trackmate movement captures. Videos can be found in the **Supplementary Information** (Supplementary Movie 1 (control) and 2 (EXO-Code)) (**c-g)** Grouped and individual plots of tracked RNA puncta distance travelled in microns, linearity of forward progression (linear speed (uM/s)/mean speed (uM/s)), distance from plasma membrane (uM), and mean velocity (uM/s). **(e)** Percentage breakdown of mobile and immobile puncta. Cellular edges were detected using a transmitted light channel and outlined prior to puncta quantification. **(h)** Categorization of puncta speed by annotated definitions of passive transport, low-speed, average-speed, and high-speed active transport. The experiment was performed in duplicate with five replicates per condition per experiment. All data presented as mean ± standard deviation. Statistical comparisons were performed using the unpaired *t*-test. \*\*\*\**p* < 0.0001, \*\*\**p* < 0.001, \*\**p* < 0.01, \**p* < 0.05. Scale bar: 10 µm

EXO-Code puncta also moved nearly twice as fast as the control puncta (**Fig. 2g)**. Given that RNA speed can be associated with different biological functions, we stratified puncta speed into “active” and “passive” transport categories, hypothesizing that the localization of EXO-Codes to sEVsrequires a faster mode of transport comparable to that of viruses (**Fig. 2h)**. Viral capsids hijack microtubule-based transport mechanisms to allow them to move at between 0.5 µM/s and >1 µM/s in order to rapidly infect target cells^35–39^. A puncta speed of 0-0.5 µM/s was regarded as passive transport, 0.5-0.75 µM/s as low-speed active transport (minimum speed of vaccinia, Ebola, and influenza viruses), 0.75-1 µM/s as average speed active transport, and >1 µM/s as high-speed active transport^37, 38^. 83.3% of the EXO-Code puncta moved at an active transport speed (>0.5 µM/s) compared to 49.2% of the control RNA; 38.3% at speeds >0.75 µM/s (comparable to kinesin-dependent vesicular transport^40^) compared to 7.7% of control RNA: and ∼7% at high-speed active transport (>1 µM/s) compared to 0% in control RNA (**Fig. 2h)**. Taken together, these data support our hypothesis that the EXO-Code is actively transported throughout the cell using similar transport mechanisms to those used by viruses and: (1) at similar subcellular speeds to viruses, (2) proximal to the plasma membrane, and (3) in a linear and directed fashion, indicating that the EXO-Code has a drastically different intracellular binding profile than that of the control sequence.

### Cellular interactome of EXO-Codes determined via proteomics and imaging

We next examined the key protein players mediating EXO-Code sorting to sEVs. Exosomal proteins are thought to mediate RNA loading into exosomes^41, 42^. Once bound to exosomal proteins, nucleic acids are packaged into invaginations from the endosomal membrane to form exosomes^3^. While Y-box protein 1, HnRNPL, and various other RNA-binding proteins have been implicated in exosomal packaging, the precise mechanisms underlying this process are poorly understood, and investigations often rely on cell-free systems or require several knockdown procedures to overcome compensatory mechanisms.^43–46^ Since several proteins may interact with EXO-Codes, we performed an RNA pull-down assay and mass spectrometry to examine whether EXO-Codes preferentially bound to subsets specific proteins. We then further probed the highest abundance proteins by confocal microscopy.

In total, 417 proteins bound to the EXO-Codes and controls. After removing proteins below the 25^th^ abundance percentile to exclude poorly binding proteins and retaining those above background, 278 proteins remained for bioinformatic analyses. Proteins that bound equally to either the control or the EXO-Code were considered non-specific binders and were not used to determine key binding proteins.

The M1 EXO-Code bound in high abundance to several classes of exosome component proteins, RNA-sorting proteins, and endocytic multi-vesicular body proteins, consistent with hijacking intracellular sorting pathways, while the control RNA sequence bound in near equal abundance to degradative proteins, cytoskeletal proteins, and ribosomal proteins despite lacking a start codon (**Fig. 3a-c**). Proteins previously identified as key exosomal miRNA-sorting components were highly bound to EXO-Code RNA, in particular SYNCRIP, HNRNA2B1, YBX1, and HNRNPM, all of which have been investigated in exosome biogenesis and cargo loading^29, 32, 33, 41, 44^. For example, SYNCRIP and HNRNA2B1 knockdown impaired exosome sorting of 55/70 endogenous murine miRNAs, while YBX1 knockdown in HEK-293T cells severely impaired miR-223 loading^33, 44^. These data indicate that the EXO-Codes are bound by a specific network of multiple proteins capable of packaging and exporting RNA via exosomes.

**Figure 3.**
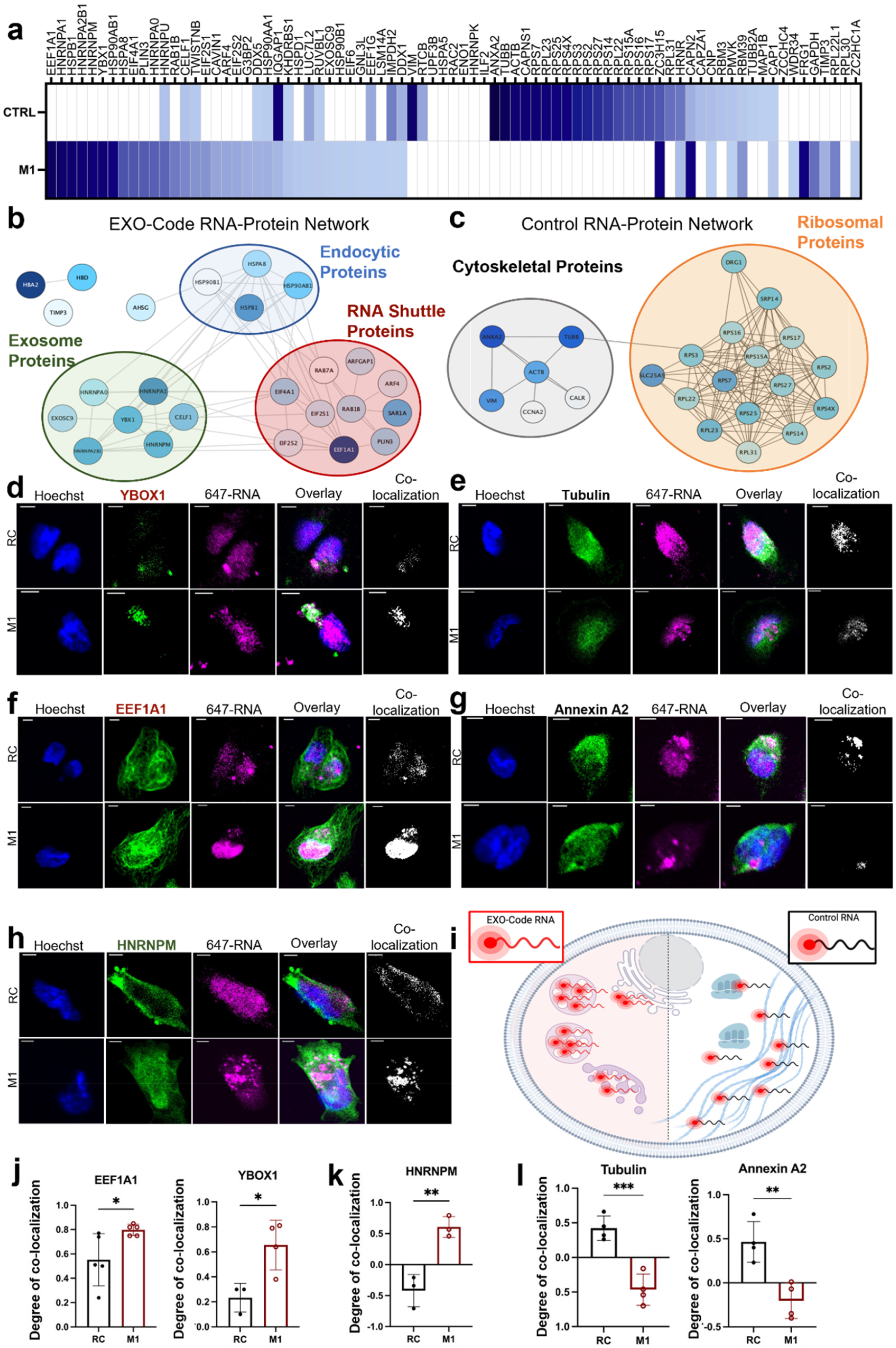
Analysis of EXO-Code binding proteins. **(a)** Semi-quantitative protein abundance values yielded from label-free mass spectrometry (Fusion Lumos) in either the control or EXO-Code RNA pulldown (n=2). (**b-c)** Cytoscape-visualized Nanostring protein interaction network for the highly abundant control and EXO-Code proteins, respectively. Interactions between proteins are visualized using Cytoscape, grouped and categorized by subcellular location. The circles representing each protein are color mapped by abundance, with dark blue indicating highly abundant proteins and white representing the least abundant proteins. **(d-h)** Confocal microscopy images assessed for co-localization with the most abundant proteins found in the pulldowns. Representative z-slices shown, and scale bars set to ten microns in each image. (**i)** Schematic of the proposed subcellular distribution as visualized by confocal microscopy, with the EXO-Code being more punctate and the control being more diffuse. **(j-l)** Pearson’s correlation coefficient (PCC) values for co-localization analysis for the sEV protein HNRNPM, RNA shuttle proteins EEF1A1, and cytoskeletal proteins tubulin and annexin a2, respectively. PCC values are defined as: −1 = negative correlation, no co-localization; 0 = no correlation, and +1 = positive correlation, high co-localization. All data presented as mean ± standard deviation. Statistical comparisons were performed using the unpaired *t*-test. \*\*\*\**p* < 0.0001, \*\*\**p* < 0.001, \*\**p* < 0.01, \**p* < 0.05.

To further investigate the potential EXO-Code sorting mechanism, confocal microscopy and co-localization analysis was performed with the highest abundance proteins within the RNA pulldown. For the EXO-Code, these studies were performed with a focus on HNRNPM, YBOX1, and EEF1A1 and, for the control, tubulin2A and annexin A2 were investigated (**Fig. 3d-l**). We hypothesized that because these proteins were among the highest abundance proteins in each respective pulldown, they would also co-localize with the fluorescent RNA after electroporation into live cells. HNRNPM and YBOX1 are both known to participate in miRNA sorting and exosome-mediated miRNA export^29, 32, 47^. While traditionally associated with transcriptional elongation, EEF1A1 expression has recently been linked both to drug and stress resistance in metastatic breast cancer, as well as active protein export toward plasma membranes, so high levels of EEF1A1 within our pulldown could further support a novel role for the protein^48, 49^. For the control sequences, we saw a high level of co-localization with tubulin and annexin A2, which did not colocalize with the EXO-Code (**Fig. 3l**). We also observed a distinctly punctuate nature of the EXO-Code puncta, which was not observed in the control sequence. Annexin A2 is thought to play a role in sequence-independent miRNA sorting into extracellular vesicles and might act as an miRNA “sponge” in autoimmune disease such as multiple-sclerosis^50, 51^. Taken together, these data provide new insights into composition of the protein networks responsible for sorting sEV-specific RNAs. With this new knowledge that EXO-Codes (1) hijack an active transport pathway and (2) bind to sEV-related miRNA sorting proteins, we next functionalized our EXO-Code to develop an RNA-based biochemical tool capable of tracking sEV intracellular dynamics and secretion.

### Rational design of an EXO-Code RNA aptamer for exosomal RNA tracking

After establishing that our EXO-Codes (1) are highly enriched into sEVs, (2) display directed movement towards the plasma membrane, and (3) co-localize with both previously annotated and novel exosome sorting proteins, we set out to develop a tool for labeling sEV RNAs that could be genetically encoded to track the entire lifespan of exosomal RNA from biogenesis to secretion.

We first created a bifunctional aptamer chimera composed of (i) our RNA EXO-Code for sEV sorting and (ii) a fluorogenic RNA Mango aptamer that when bound to thiazole orange-1 (TO-1), or thiazole orange-3 (TO-3) increased its fluorescence thousands of times^12, 13, 52^. To assess the stability of the RNA Mango-TO-1 complexes, we peformed an inverse cell uptake experiment, where the permeability of the TO-1 alone was compared with that of TO-1 complexed with RNA. TO-1 alone was cell permeable, but not when complexed with the RNA Mango. Therefore, TO-1 stably intercalates into the RNA aptamer and neutralizes uptake of free dye molecules (**Supplementary Fig. 3**).

To determine structure-activity relationships, we first designed several EXO-Code/RNA Mango fusion constructs, assessed their secondary structures and thermodynamic stabilities and assessed if fusing of the EXO-Code to the RNA Mango affected TO-3 binding and fluorescence intensity (**Fig. 4a-c, Supplementary Table 1**).

**Figure 4.**
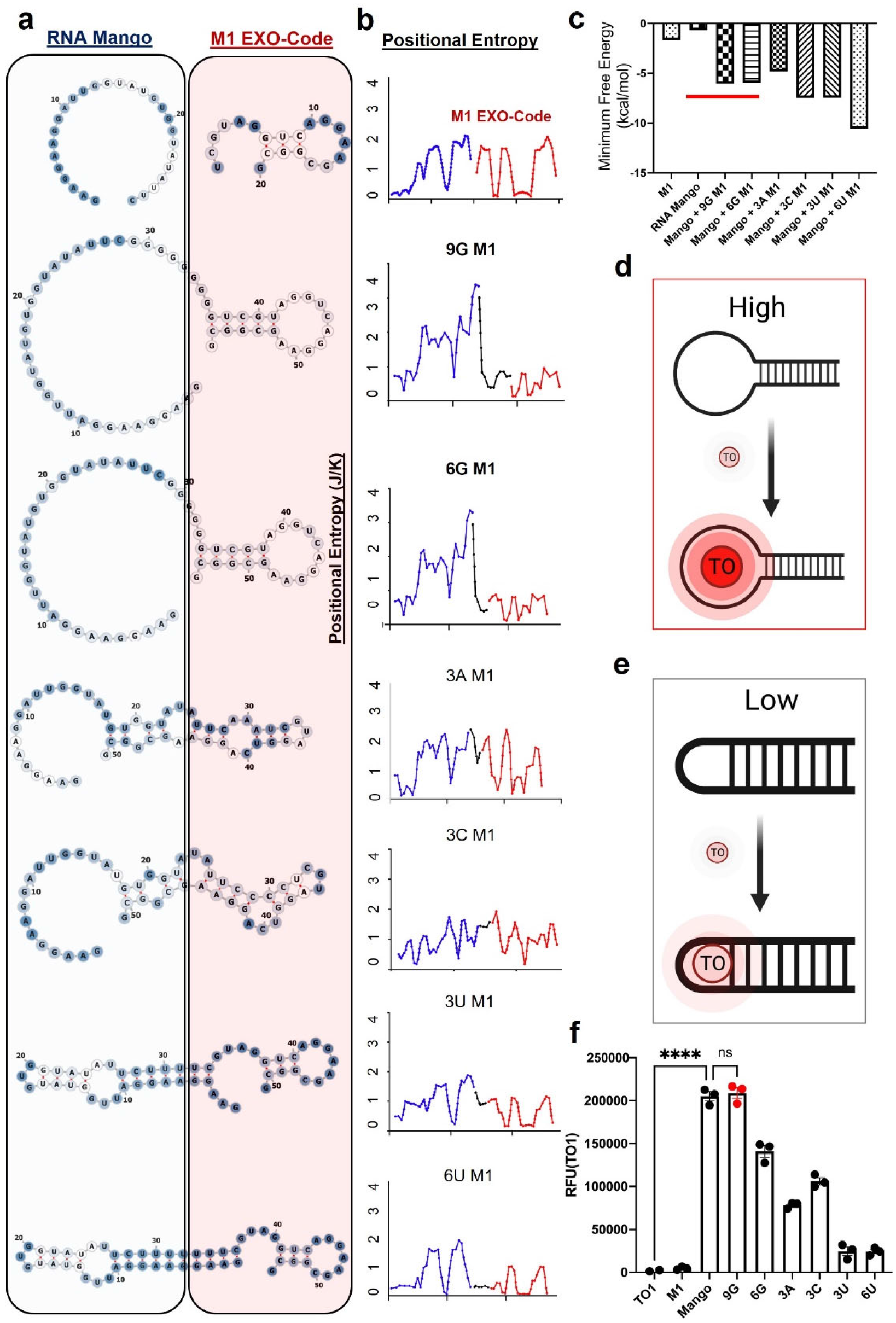
Secondary structure predictions of EXO-Probe aptamers show altered fluorophore binding efficiency. **(a)** Schematics depicting the *in silico* determined secondary structures of RNA EXO-Probes color-coded by positional entropy, with dark blue indicating a more rigid structure (low positional entropy) and white indicating more flexibility (high positional entropy). (**b)** Positional entropy plotted as chimera components as blue (Mango), black (linker), and red (EXO-Code). (**c)** Minimum free energy predictions of secondary RNA structures shown in panel (a). The red bar across the RNA Mango, 9G M1 and 6G M1 structures indicate those with the highest fluorescence in panel (**f).** (**d,e**) Schematic depicting fluorescence enhancements when secondary structure of the RNA Mango is maintained within the final EXO-Probe structure. (**f**) *In vitro* assessment of fluorophore binding efficiency as compared to the RNA Mango aptamer alone. Experiment was conducted in triplicate, and a representative assay is presented. All data presented as mean ± standard deviation. Statistical comparisons were performed using a one-way ANOVA \*\*\*\**p* < 0.0001, \*\*\**p* < 0.001, \*\**p* < 0.01, \**p* < 0.05.

Since the secondary and thus tertiary structure of RNA can be extremely diverse and dictate biological function, we assessed several structures with different linker lengths to ensure that (1) our EXO-Code would remain active as a molecular zip code and (2) that RNA Mango’s fluorophore-binding capability would not be hindered as visualized in **Fig. 4d,e**^53–56^. We hypothesized that the top performing EXO-Probe would have a stable secondary structure that would not markedly change with addition of RNA Mango compared to their free counterparts. Six structures were designed and synthesized with increasing polynucleotide linker lengths (**Fig. 4a**). Conjugation of the EXO-Code to RNA Mango increased the predicted thermodynamic stability of the constructs ∼8-fold compared to their individual components and in some cases, dramatically altered the entropic profile of species (**Fig. 4b,c**). Structures that maintained both the RNA Mango and the RNA EXO-Code secondary structure (9G, 6G) showed the highest fluorescence, as they maintained the required G-quadruplex necessary for TO fluorophore binding. Two structures that partially maintained the RNA Mango secondary structure and the EXO-code (**Fig. 4a, 4c**) showed a 55% decrease in fluorescence while 3U and 9U with predicted abrogation and “zipping up” of the RNA Mango secondary structure had only 12% of the fluorescence intensity of the original RNA Mango structure when incubated with TO-3 (**Fig. 4f**).

### EXO-Probes as a genetically encodable exosome imaging agent

The 9G-M1 conjugate exhibited approximately 15,000-fold and 10-fold greater relative fluorescence intensity than that of the fluorophore or EXO-Code alone, respectively and did not decrease significantly compared to RNA Mango alone **(Fig. 4f**). Since 9G-M1 maintained nearly 100% of the secondary structure and entropic profile of RNA Mango, we next determined whether this promising candidate also maintained enrichment of the bifunctional EXO-Code/RNA Mango chimera into exosomes (**Fig. 5**). Upon electroporation into MDA-MB-231 cells, the 9G-M1 aptamer showed ∼350-fold greater enrichment into sEVs than the control or the Mango aptamer alone (**Fig. 5b**).

**Figure 5.**
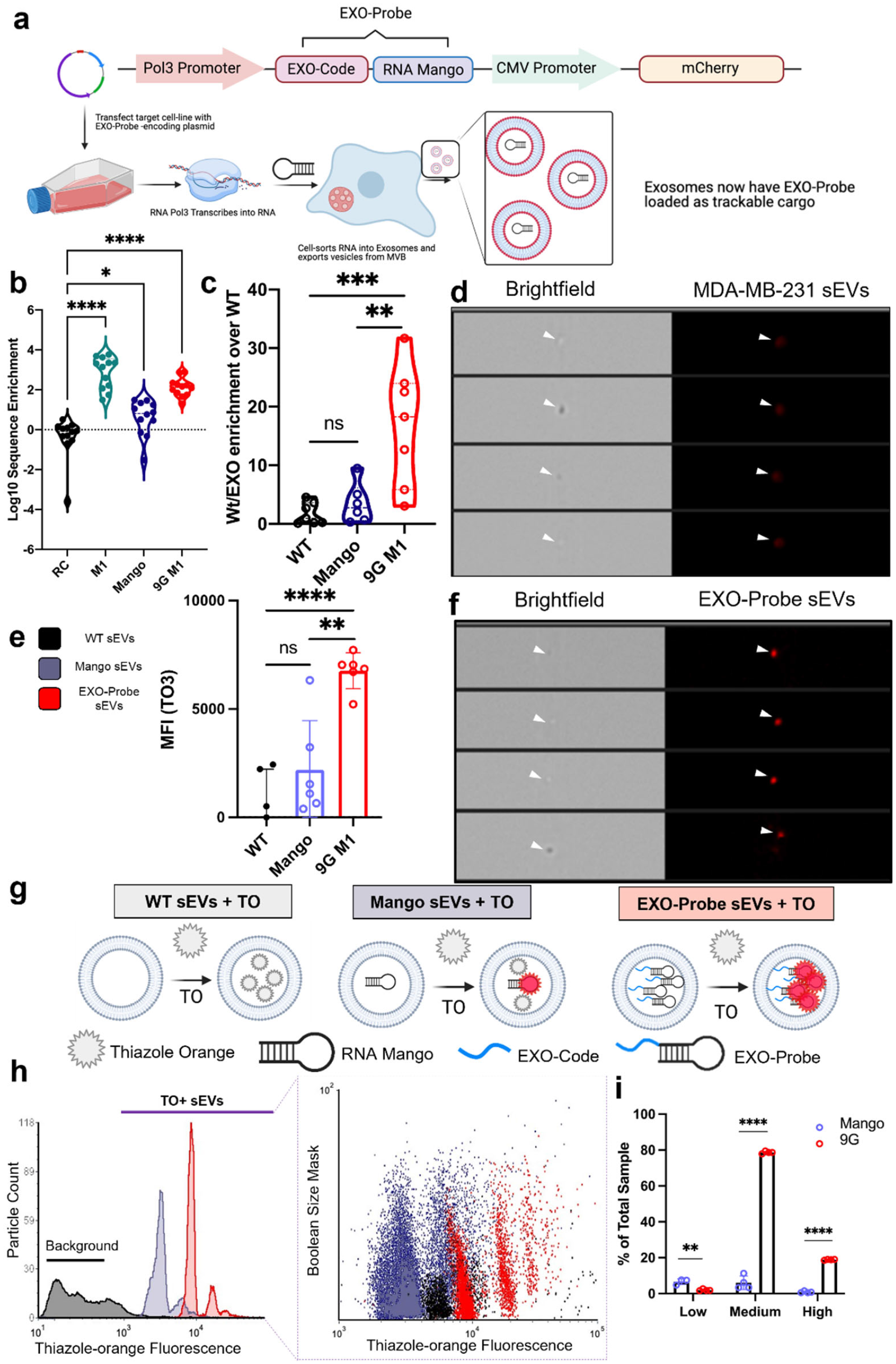
Functional assessment of the EXO-Probe aptamer. **(a)** Plasmid map of the shRNA-expressing plasmid used to generate genetically-encoded EXO-Probe aptamers, highlighted with black brackets as a combination of the EXO-Code and RNA Mango. The EXO-Probe construct was placed under the control of the RNA polymerase III (Pol III) promoter, and the mCherry reporter protein was under control of the CMV promoter. The lower half is a schematic depicting the proposed route of genetically encoded EXO-Probe sEV enrichment and categorization. The plasmid is transfected into the target cell line, wherein RNA Pol III transcribes the RNA, which is then subsequently packaged into sEVs. **(b)** Quantitative RT-PCR enrichment comparison of RNA EXO-Probes and EXO-Codes compared with random control and RNA Mango aptamer. **(c)** Quantitative RT-PCR data of genetically-encoded EXO-Probe enrichment after plasmid transfection. Plasmids were transfected into cells and grown in sEV-depleted medium, after which the sEVs were harvested and RNA extracted to obtain data in panel (**b-c). (d-f)** Single particle sEV flow cytometry images highlighting the contrast between EXO-Probe loaded sEVs and dye-labeled sEVs and quantification plot represented as mean +/-standard deviation. **(g)** Schematic representing the proposed system of EXO-Probe loading and labeling at the sEV level. **(h)** Flow cytometry histogram of data in panels (d-f) highlighting gating methodology. Instrument debris and negative sEVs were gated out based on SSC and image quality. Dot plot gated based on multi-channel Boolean size masks of TO+ sEVs directly compared with EXO-Probe loaded sEVs. (**i)** Striation of three distinct populations observed in the sEV flow cytometry dot plot and histograms. Data is presented as the percent of the total sample analyzed, +/-standard deviation. Experiments (**b-i)** were completed in quadruplicate, and representative experiments are shown. Data are presented as mean ± standard deviation. Statistical comparisons were performed using a one-way ANOVA \*\*\*\**p* < 0.0001, \*\*\**p* < 0.001, \*\**p* < 0.01, \**p* < 0.05. Statistical comparisons for panel i were done using a student’s t-test to determine significance between the means with \*\*\*\**p* < 0.0001, \*\*\**p* < 0.001, \*\**p* < 0.01, \**p* < 0.05.

Having confirmed sEV enrichment of the 9G-M1 RNA (**Fig. 5b**), we next genetically encoded the system, as there is currently no genetically encoded material capable of both enriching into sEVs whilst emitting fluorescence upon fluorophore binding. We therefore constructed a gene encoding the M1 EXO-Code fused to the RNA Mango aptamer and cloned it into an shRNA-producing lentiviral expression vector (**Fig. 5a, Supplementary Table 2**). Successful transfection was assessed via the co-expressed mCherry reporter protein by flow cytometry after which RNA from the sEVs of transfected cells was assessed for M1 EXO-Code/RNA Mango (**Supplementary Fig. 4**). There was 20-fold enrichment of 9G-M1, herein referred to as an “EXO-Probe”, within the sEVs of MDA-MB-231 cells (**Fig. 5c**). Using hybrid imaging single-particle flow cytometry, we optimized instrument settings and calibration for sEVs using polystyrene 100 nm size standard beads (**Supplementary Fig. 5**) which allowed us to quantify the fluorescence and capture images of sEVs for subsequent experiments. With this sEV-specific hybrid flow cytometry protocol, we detected ∼8-fold enhancement in sEV fluorescence after incubation with the thiazole orange dye (**Fig. 5e, g-h**). The sEVs had three distinct populations of varying fluorescence, defined as “low”, “medium”, and “high” (**Fig. 5i**). sEVs loaded with the EXO-Probes had significantly higher numbers of sEVs displaying medium and high fluorescence compared to the unloaded, TO-3 stained sEVs. This phenomenon has been observed in previously published studies, and is thought to be due to cellular secretion of distinct subpopulations of sEVs of varying RNA and protein densities^57–59^.

Visualization via the Amnis Imagestream (**Fig. 5d, f**) showed EXO-Probe-loaded sEVs displaying significantly brighter fluorescence than that of unloaded, TO-3 stained sEVs. Therefore, we concluded that our RNA was not only successfully transcribed but also maintained its dual functionality as a molecular zip code and fluorogenic RNA aptamer.

### Genetically encoded EXO-Probes display spatial proximity to sEV RNA sorting proteins

To verify that our genetically encoded EXO-Probe was indeed sorted via intracellular pathways, we re-performed live cell imaging to track RNA puncta within transfected MDA-MB-231 cells. Distinct intracellular puncta were observed in both the RNA Mango and the EXO-Probe plasmid transfected groups, indicating that the free dye could permeate and bind RNA aptamers (**Fig. 6a**). The EXO-Probe puncta had on average twice the mean straight line speed than the encoded RNA Mango controls, indicating a higher degree of linear and directed movement (**Fig. 6b, Supplementary Movies 3 and 4**). This was further supported by the EXO-Probe possessing twice the linearity of forward progression compared to the RNA Mango control **(Fig. 6c**). On average, the EXO-Probes also possessed a lower mean directional change rate, further indicating that the puncta were not moving randomly but in a more directed manner than the RNA Mango alone (**Fig. 6d**).

**Figure 6.**
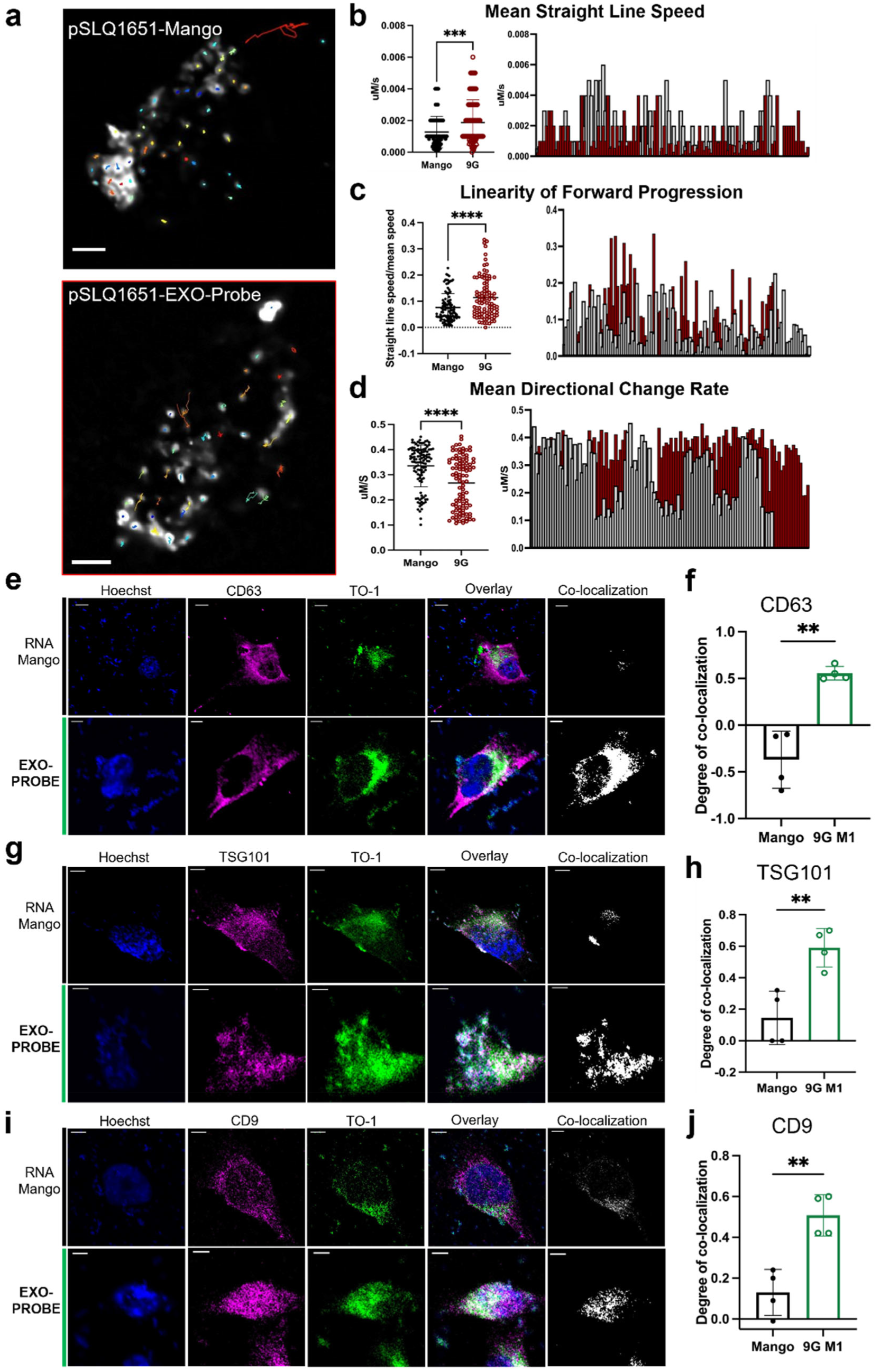
Genetically encoded EXO-Probe subcellular movement and distribution. **(a)** Live cell stills depicting TO-1(+) Trackmate particle movement within pSLQ1651-transfected RNA Mango (top) and EXO-Probe 9G-M1 (bottom) cells. (**b-d)** The Trackmate parameters depict straight line speed, mean directional change rate, and linearity of forward progression, shown as individual puncta. Experiments a-d were performed in duplicate, and representative video stills are shown. Full videos can be found in the **Supplementary Movie 3 (Mango control) and 4 (EXO-Probe). (e-j)** Confocal microscopy images of genetically encoded RNA Mango and RNA EXO-Probes co-localized with sEV-specific tetraspanins CD63, TSG101, and CD9, and quantification of co-localization via Pearson’s correlation coefficients (PCC). PCC values are defined as: −1 = negative correlation, no co-localization; 0 = no correlation, and +1 = positive correlation, high colocalization. Scale bar = 15 µm. Experiments were conducted in triplicate, and all data are presented as mean ± standard deviation. Statistical comparisons were performed using the unpaired *t*-test. \*\*\*\**p* < 0.0001, \*\*\**p* < 0.001, \*\**p* < 0.01, \**p* < 0.05.

The genetically encoded EXO-Probes also co-localized with the sEV-specific endosomal marker CD63, while RNA Mango did not (negative degree of co-localization)^60^ (**Fig. 6e,f**). CD63 has been used to design novel and sophisticated sEV membrane labeling systems, so it was highly encouraging to see that our EXO-Probes co-localized with this marker^25, 26^. Furthermore, the EXO-Probe was significantly more co-localized with two additional sEV markers, TSG101 and CD9, than the non-targeted RNA Mango (**Fig. 6g-j**). These data, when combined with qPCR and sEV flow cytometry data, support our hypothesis that the EXO-Probe (1) maintains the dual function of an sEV-specific molecular zip code and fluorogenic RNA aptamer, (2) hijacks similar intracellular sorting pathways to the EXO-Code alone, and (3) is capable of being genetically encoded into cells while maintaining these functions. With this new knowledge, we then sought to implement our EXO-Probe as a simple means to quantify sEV secretion.

### EXO-Probes allow relative sEV quantification via genetic encoding and fluorophore pulsing

To establish an sEV biochemical toolkit, we hypothesized that our EXO-Probe would be detectable via fluorophore binding and pulsing within uncentrifuged cell culture medium. Having verified that the EXO-Probe has a greater affinity for the TO-1 fluorophore than controls after cellular loading into sEVs, we developed a semi-quantitative assay that exploits the RNA-TO-1 interaction to label and quantify MDA-MB-231 sEVs. This represents one of the few assays capable of labeling RNA-loaded sEVs without ultracentrifugation or other pre-purification methods and which does not require membrane modifications of the sEVs. As shown by NTA and single particle flow cytometry data, EXO-Probe loading into sEVs and incubation with the fluorophore did not significantly alter their size significantly from that of sEVs from unmodified MDA, or sEVs labeled with the TO-1 fluorophore. This new assay will allow investigators to easily detect and quantify sEVs without the need for ultracentrifugation. To test this hypothesis, we electroporated our EXO-Probe into MDA-MB-231, harvested conditioned media, and incubated it with the TO-1 fluorophore (**Fig. 7a,b**). Exosomes secreted into the cell culture medium were quantifiable and showed a strong correlation with exosome number as quantified by NTA after differential centrifugation (correlation coefficient >0.94). Wild-type (WT) conditioned medium and sEVs had fluorescence above exosome-free media controls. This could be due to the TO dye partially intercalating into other nucleic acids within the bulk EV population^12^. The increase in the EXO-Probe conditioned medium volume was highly correlated with an increase in overall fluorescence, as evidenced by a correlation coefficient of >0.95 for the conditioned medium and >0.99 for the sEVs, respectively (**Fig. 7c**).

**Figure 7.**
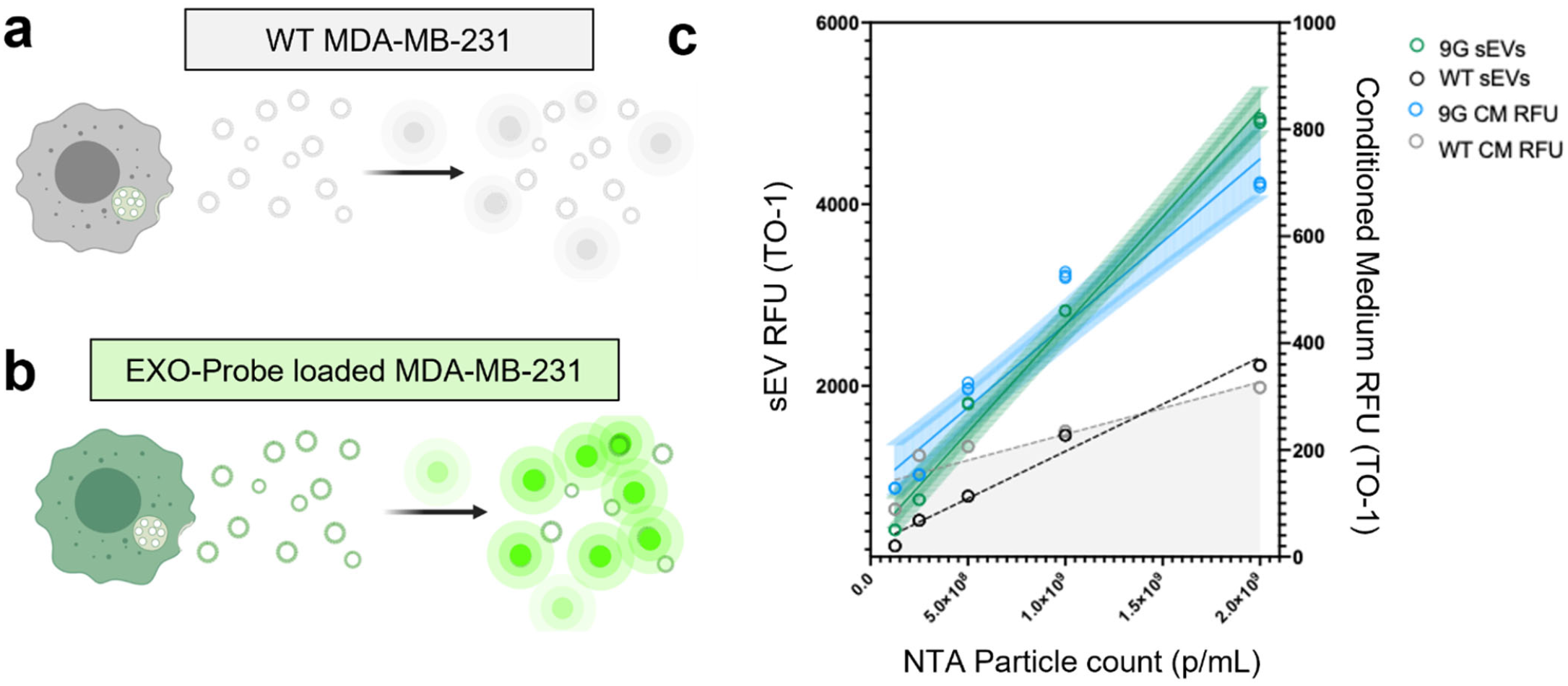
The EXO-Probe as an RNA tool for sEV quantification. (**a-b**) Schematic depicting that (**a**) without the EXO-Probe aptamer, unmodified or wild-type (WT) sEVs will show little fluorescence when incubated with an RNA dye. (**b**) When cells are loaded with the EXO-Probe, the resulting sEVs will exhibit high fluorescence. (**c**) Fluorescence data presented as the relationship between sEV RFU and conditioned medium (CM) RFU after incubation with TO-1 and compared to NTA particle counting. The blue and green lines indicate the RFU values of the EXO-Probe loaded sEVs and CM, respectively. The dashed gray and black lines represent the unmodified or wild-type (WT) CM and sEVs, respectively. All data are presented as individual data points with a simple linear regression fit to the points. The correlation coefficients are 0.99, 0.94, 0.98, and 0.81 for the EXO-Probe sEVs, CM, WT sEVs, and WT CM, respectively. 9G CM and 9G sEVs are shown with 95% confidence intervals. Experiments were conducted in triplicate, and representative data are shown. All data are presented as mean ± standard deviation. Linear regression model fitted using GraphPad Prism. Statistical considerations of slope significantly greater than zero shown as \*\*\*\**p* < 0.0001, \*\*\**p* < 0.001, \*\**p* < 0.01, \**p* < 0.05.

## Discussion

There is increasing interest in understanding sEV biogenesis, release, and subsequent uptake into recipient cells. Several strategies have been developed for this purpose, such as pHlourin, a GFP mimic tagged to the tetraspanin CD63, which has been used to track exosome secretion from migratory cells and cellular uptake and endocytosis by recipient fibro-carcinoma cells. While this and other exosome membrane labels are highly useful, they label the sEV membrane and do not label the sEV RNA cargo. Also, while tetraspanins CD9, CD63, and CD81 are key players in exosome biogenesis, not all subsets of sEVs and exosomes display them so unique subsets may not be detected with sEV membrane labeling techniques^16, 57, 59^. Furthermore, the overexpression of tetraspanins can alter both sEV secretion from cells and their mechanism of uptake, thus altering their downstream effects^27^.

Until now, there has not been a method to label the internal RNA cargo of sEVs via genetic encoding to track exosomal RNA sorting and biogenesis from formation to cellular uptake and release into the cytoplasm. Other genetically-encoded materials used for monitoring sEV RNA transfer, such as the CROSS-FIRE system, are not targeted toward EV-mediated RNA sorting^61^. We believe that our genetically-encoded, sEV targeted system could be used in conjunction with this type of labeling to better understand sEV RNA movement at the single-cell level. Specific methods and dyes for RNA labeling are limited, thus making tracking of sEV RNA challenging. This challenge is further complicated by each RNA species being present at extremely low copy numbers. To achieve sufficient sensitivity, we established a new screening method to identify highly enriched RNA sequences in sEVs. These sequences were >1000-fold greater enriched into sEVs over control sequences and published motifs. Moreover, the method was applicable to other cell types such as mesenchymal stem cells and others (**Supplementary Figure 1**).

We also designed a genetically encoded exosomal RNA tracking tool by fusing our EXO-Code to an RNA aptamer, termed RNA Mango, which specifically binds to thiazole orange (TO). The top-performing structure both in terms of RNA enrichment and fluorophore binding, 9G-M1, was predicted to maintain 100% of the RNA Mango secondary structure and >70% of the unconjugated EXO-Code structure. We verified that 9G-M1 maintained dual functionality as an EXO-Code and a TO-binding aptamer through quantification of intracellular trafficking, qPCR, and single particle flow cytometry. These data provide strong evidence that the secondary structure of dual-functional RNA must be taken into consideration before designing RNA aptamers.

Specifically, we examined the characteristics of individual RNA puncta, monitoring their kinetic parameters and intracellular sequestration. We observed a clear subpopulation of highly mobile EXO-Code RNA molecules moving at comparable speed linearity to RNA viral capsids. These puncta were also nearly all localized close to the plasma membrane rather than immobilized in intracellular pockets. These findings provide strong evidence for the directed movement of EXO-Code RNA. Intracellular RNAs without transport or localization signals remain subject to the forces of diffusion and random movement to reach the perinuclear region or a ribosome in the case of siRNA or mRNA, respectively^62–64^. Thus, only a small fraction reaches the multivesicular bodies for packaging into sEVs. Taken together, these data demonstrate that the EXO-Code RNA puncta utilize the intracellular sorting machinery to be directed to distinct subcellular compartments and together with RNA-protein pulldowns, that EXO-Codes use an active transport mechanism into sEVs rather than passive diffusion, which only transports <0.1% of non-targeted RNA into sEVs.

We quantified EXO-Code-interacting proteins using pulldown and mass spectrometry techniques and identified distinct protein families that might mediate exosome miRNA cargo loading including RNA shuttle proteins, endocytosis proteins, and exosomal proteins. Of these proteins, several are known to bind to RNA motifs present in the EXO-Code sequence. YBOX1 specifically associates with short, four nucleotide RNA motifs containing guanine and uracil. Recent work analyzing heterogenous ribonuclear proteins found that HNRNPM, HSP90AB1, and HNRNA2B1 effectively shuttle small RNAs into exosomes, and may themselves act to protect sEV RNA^41^. We observed that these proteins not only bound the EXO-Codes but bound to a significantly higher degree than to non-specific RNA.

We also found that EEF1A1 was a specific EXO-Code binding partner as indicated by our proteomic and confocal microscopy data. While EEF1A1 is traditionally thought of as a mediator of tRNA binding to ribosomes, it is also known to promote transcription in T helper cells and localize with proteins near cellular membranes^65^. Our data suggest a potentially novel role for EEF1A1 in exosome miRNA cargo packaging, a hypothesis that requires testing. In contrast, the control RNA sequence bound more strongly to cytoskeletal proteins and ribosomal proteins. One hypothesis is that by binding primarily to cytoskeletal proteins, such as actin, tubulin, annexin, and ribosomal proteins, the RNA control sequence becomes trapped within the cytoplasm. Consistent with this, a large proportion of RNA control sequence was immobile, with a small fraction remaining diffuse within the cytoplasm.

Our screening and bioinformatic secondary structure prediction allow the control of sorting and movement of RNA molecules. Our EXO-Probe can also be genetically encoded into cells and, in the future, into transgenic mice or xenografts, allowing *in vivo* exosome tracking and characterization. Unlike protein-based fluorophores, labeling exosomal RNA through RNA aptamers allows the easy exchange of fluorophores therefore providing significantly more flexibility in terms of multi-color imaging without the need for time-consuming recloning of optimized genetically-engineered constructs. In our case, the RNA Mango can bind both to TO-1 (em/ex 480/530 nm) and TO-3 (em/ex 635/658 nm), but we expect the addition of new dyes and fluorophores as the field of RNA aptamer fluorophores matures.

We were further able to verify that our genetically-encoded EXO-Probes utilized endocytic pathways, as they interacted closely with the sEV secretion-specific proteins CD63, Tsg101, and CD9. We specifically interrogated the subcellular movement of our EXO-Probes through live cell imaging and found that the EXO-Probes indeed moved faster and in a more linear fashion than that of the non-targeted RNA Mango sequence alone. This led us to test an innovative application of this technology, namely using our RNA aptamer to assess relative sEV quantity from conditioned medium alone without the need for ultracentrifugation or other purification and isolation techniques. Furthermore, we developed an sEV labeling assay that worked with either conditioned medium or ultracentrifuged sEVs.

In summary, here we report the first exosomal RNA labeling toolkit that can be genetically encoded into cells. This technology will be useful for studying mechanisms of RNA trafficking to sEVs and the underlying cellular sorting decisions. Furthermore, it will enhance our understanding of the role of sEVs in health and disease across a broad range of disciplines including biomedicine, immunology, neurobiology, cancer biology, and more. In the future we seek to further optimize our EXO-Probe technology and develop biochemical toolkits capable of tracking the movement and deposition of sEV cargo.

## Author Contributions

Research idea, experimental study concept and design, J.N., E.B. Conceptualization and design of EXO-Code screening J.N. and S.F.. Acquisition of data, E.B., S.F., N.J., J.W., A.D.B, D.K. EXO-Code screening and characterization, S.F., J.W., E.B., N.J. Live-cell imaging, D.K, M.S.I. Data analysis, E.B, J.N. S.F. J.W. D.K. RNA Seq analysis, proteomic data analysis, E.B. Confocal microscopy, flow cytometry, molecular cloning, RNA structure design, EXO-Probe secretion, E.B. Statistical analysis, E.B. Drafting and writing of manuscript, J.N. and E.B. Study supervision, J.N. Obtained funding, J.N.

## Acknowledgements

We acknowledge funding through the National Institutes of Health (R01EB023262) and the National Science Foundation DMR 2000256. Parts of figures 1, 4, 5, and 7 were created using Biorender.com. Images were acquired at the UNC Neuroscience Microscopy Core (RRID:SCR_019060), which is supported, in part, by funding from the NIH-NINDS Neuroscience Center Support Grant P30 NS045892 and the NIH-NICHD Intellectual and Developmental Disabilities Research Center Support Grant P50 HD103573. This work has been made possible in part by grant number 2019-198107 to M.S.I. from the Chan Zuckerberg Initiative DAF, an advised fund of Silicon Valley Community Foundation.

## Financial Competing Interests

JN, SF, and EB are inventors on the patent applications of the EXO-Code technology evaluated in this paper that have been licensed to Exopharm and have received royalties. These relationships have been disclosed to and are under management by UNC-Chapel Hill.

## Methods

### Cell culture

All cells were maintained at 37°C, 5% CO_2_ and passaged upon confluency prior to POSTAL screening as per established cell culture protocols. MDA-MB-231 cells and MSCs were cultured in high-glucose DMEM + L-glutamine (Gibco, Thermo Fisher Scientific, Waltham, MA) supplemented with 10% fetal bovine serum (Genesee Scientific, San Diego, CA), or MESENPRO RS Medium (Life Technologies, Carlsbad, CA), respectively. Cells were detached using 1 mM EDTA in PBS (Gibco), and 20 µg of an RNA library (TriLink BioTechnologies, San Diego, CA) of 10^12^ random 20-nucleotide sequences was electroporated into a range of three to six million cells using the NEON Transfection System (Invitrogen, Carlsbad, CA). MDA-MB-231 and MSCs were electroporated using the following parameters: 1400 V, 10 ms, 4 pulses, and 990 V, 40 ms, 1 pulse, respectively. After electroporation, cells were suspended in 500 µL DMEM and were subsequently spun at 500 x g for 5 min. Medium containing excess RNA was removed, and this process was repeated five times to remove any RNA from outside of the cells. Cells were then resuspended and plated into flasks containing EXO-depleted medium (high glucose DMEM + L-glutamine + 10% EXO-depleted FBS, preparation of which described here^66^). Cells were cultured for 96 h to ensure sufficient accumulation of sEVs.

### Vesicle isolation and RNA extraction

Medium was collected from the cells and spun at 2,500 x g to remove live cells and large debris. The supernatant was then combined with one-half volume of Total Exosome Isolation Reagent (Invitrogen) and collected using the manufacturer’s protocol. Quantity and size distribution of exosome samples were determined using Nanosite tracking analysis (NTA). Harvested sEVs were resuspended in PBS prior to analysis on the NTA. 1 µL RNAse A/T1 (Life Technologies) per 100 µL exosome sample was added to digest RNA not within sEVs. The samples were then incubated at 37°C for 30 min. RNA was extracted from the sEVs using the miRNeasy kit (Qiagen, Hilden, Germany) following the manufacturer’s protocol.

### PCR amplification of RNA library

Since the RNA pool contained both endogenous exosomal RNA as well as that from within the selection rounds, RNA was amplified using 2 µL of 200 µM 3’ primer specific for the adapter region of the library sequences. The PCR reaction contained: RNA, 5X superscript II buffer (Thermo Fisher Scientific), DTT (Thermo Fisher Scientific), 2.5 µL of 4 mM dNTPs (New England Biolabs, Ipswich, MA), and SuperScript II RT Enzyme (Thermo Fisher Scientific). This reaction mixture amplified only RNA containing the adapter sequences, and this cDNA was then transcribed to DNA using the following steps. 10 µL of cDNA was combined with nuclease-free water, 4 mM dNTPs, standard Taq buffer (New England Biolabs), 2 µL of 20 µM of each primer, and 0.5 µL of Taq polymerase (New England Biolabs). The forward primer in this reaction contained the T7 RNA promoter sequence. These sequences were then amplified 20-25x as follows: 30s at 90°C, 30 s at 56°C, and 45 s at 72°C. DNA quality was assessed using a 2% agarose gel and purified using the GeneJet PCR kit (Thermo Fisher Scientific). DNA was then eluted to a concentration of between 25-100 ng/µL and amplified using the T7 MegaShortScript RNA synthesis kit (Ambion, Austin, TX). The transcribed RNA was combined with Novex Loading buffer (Thermo Fisher Scientific) and run at 180 V for 50 min on a 10% acrylamide TBE-Urea Novex gel (Thermo Fisher Scientific). Gels were stained with 2 µg/mL ethidium bromide (Amresco, Solon, OH) in 1 X TBE and rocked for 15 min. Gels were then washed in TBE buffer for an additional 15 min with rocking. RNA was visualized with the GeneDirex LED Transilluminator, and appropriate bands were excised. Excised bands were added to 100 µL of extraction buffer (0.5 mM ammonium acetate, 1 mM EDTA, 0.2% SDS in 1X TBE) and flash frozen in liquid nitrogen. The resulting pellet was incubated with agitation for 1 h at 37°C, vortexed, and kept on ice. An additional 100 µL of extraction buffer was added to the mixture, which was subsequently centrifuged to remove smaller gel pieces. The collected extraction buffer was combined with 1.5 volumes of cold 100% ethanol, and the mixture spun on an RNA-cleanup column (Qiagen; miRNeasy). This RNA was then recovered and re-electroporated into cells at the reduced quantity of 7.5 µg. The tenth and final round of POSTAL selection was further decreased to 1 µg of RNA.

### Next-generation sequencing of isolated RNA

The harvested RNA was sequenced using an Illumina MiSeq200 at the Genomics and Bioinformatics Core at the University at Buffalo. For both MDA-MB-231 and MSCs, 10 rounds of sequencing were performed with the Illumina platform reverse-transcription technology. The following primers were used for amplification: FP: TCGTCGGCAGCGTCAGATGTGTATAAGAGACAGTAGGGAAGAGAAGGACATATGA; RP: GTCTCGTGGGCTCGGAGATGTGTATAAGAGACAGTCAGTGGTCATGTACTAGTCAA. The sequence reads from this procedure were then obtained and analyzed as detailed below.

### Bioinformatic processing of sequence reads

Raw data files were trimmed of sequencing adapters and primers using the Python package Cutadapt v.2.1. The removed adapters were also quantified against the total number of sequences to ensure that trimming had worked properly, as >98% of sequences were trimmed in each set of sequences. Reads less than ten nucleotides were removed from downstream analysis. The trimmed datafiles were then analyzed using the Perl-based toolkit FASTAptamer developed by the Burke Laboratory^67^. Sequences were first counted to determine the abundance in the respective population file. Sequence abundance was normalized by reads per million (RPM) and sorted by decreasing abundance. The sequence distributions from each selection round were then compared against each other via their respective log_2_ RPM (RPMy/RPMx). The top 400 sequences from the final round of sequencing were then converted into FASTA format and loaded into the <u>m</u>ultple <u>e</u>m for <u>m</u>otif <u>e</u>licitation database (MEME) and analyzed under classic discovery mode with a minimum sequence length of 6 nucleotides and a maximum sequence length of 20 nucleotides. The top 10 sequences from MDA-MB-231 were used for endpoint analysis and are henceforth referred to as “EXO-Codes”.

### Endpoint exosome enrichment of top performing sequences

In order to re-assess the robustness of the POSTAL selection procedure, the top performing EXO-codes were re-introduced into cells via electroporation of 10 µg of EXO-code RNA (IDT Technologies, Coralville, IA) per 330,000 MDA-MB-231 or 66,000 MSCs. Electroporated cells were centrifuged at 500 x g for 10 min and the medium containing excess RNA removed. Cell pellets were then resuspended in EXO-depleted media and plated in triplicate in a 12- or 24-well culture dish with appropriate volumes of EXO-free media for MDA-MB-231 cells or MSCs, respectively. Cells were then cultured for 72 h and sEVs harvested from the media by differential ultracentrifugation: 500 x g for 10 min to remove cells; 2,000 x g for 15 min to remove larger debris; 10,000 x g for 30 min to remove protein aggregates; and 100,000 x g for 70 min to pellet sEVs. All centrifugation steps were performed at 4°C and the supernatant harvested in all steps prior to the 100,000 x g spin. Exosome samples were harvested in 20 nm filtered dPBS and subsequently quantified and characterized using the Malvern Zetasizer prior to RNA isolation. Exosome samples were treated with 1 µL RNAse A/T1 per 100 µL sample for 30 min at 37°C and lysed in 650 µL of QIAzol (Qiagen). To monitor the RNA recovery percentage as a function of the isolation procedure, 1 ng of *C. elegans* RNA spike-ins 39 and 54 were added to the lysed sEVs as an experimental quality control. Exosomal miRNA was isolated using the Qiagen miRNeasy kit according to manufacturer’s protocol. cDNA was synthesized using the Quantabio Qscript miRNA cDNA synthesis kit (Qiagen) following the manufacturer’s instructions. qPCR standard curves of each EXO-Code and various controls were generated using serial dilutions of cDNA originating from 1 µg stock RNA. Samples were aliquoted into 10 µL qPCR reaction volumes consisting of 0.2 µL EXO-Code-specific or spike-in-specific primer (IDT), 0.2 µL universal reverse primer (QuantaBio, Beverly, MA), 5 µL SYBR green universal reaction mix (Bio-Rad, Hercules, MA), and 2.5 µL sample cDNA or 1 µL standard cDNA. Volumes were brought to 10 µL with nuclease-free water. The 2-step cycling protocol of 95 °C (2 minutes) followed by 40 cycles of 95°C (5 seconds), 60°C (30 seconds, collect fluorescence) data was used for this analysis. Exosome enrichment was determined using the respective standard curves for each EXO-Code as well as normalization by the number of sEVs quantified via NTA. This yielded a value of ng RNA per exosome. These results were then compared to that of the random control sequence and plotted as weight (WT) per exosome (EXO) enrichment over random control (RC).

### Point mutations to determine sequence specificity

EXO-Code sequences with point mutations in key motif and structural positions were assessed for exosome enrichment using the same protocol as the endpoint enrichment analysis. Point mutated RNA (IDT) was generated by swapping a single nucleotide with the opposite class (purine to pyrimidine or vice versa) and the opposing base pair to provide a comprehensive knock-out of any sequence-specific RNA sorting pathways.

### Live cell imaging of Alexa-488 tagged EXO-codes

Target cell-lines were trypsinized and counted prior to electroporation with both the appropriate amount of RNA and the proper NEON transfection system settings, as described above. The RNA used for live-cell imaging was tagged with an Alexa-488 fluorophore (IDT). 500,000 cells were plated on Nunc glass-bottomed dishes (Life Technologies) and incubated for 12 h at 37°C prior to imaging. Immediately prior to imaging, cells were stained with SYTO Deep Red nuclear dye (excitation/emission maxima 652/669 nm, Thermo Fisher Scientific). SYTO Deep Red was diluted 1:2000 in high-glucose Phenol-Free DMEM (Thermo Fisher Scientific). Culture medium was removed from cells, and cells were incubated in SYTO Deep Red-containing medium for 30 min. After incubation, SYTO Deep Red solution was aspirated, cells were washed twice with sterile filtered PBS, and cell culture medium was replaced. Images were acquired on an inverted Olympus FV3000RS confocal equipped with a Tokai Hit stage-top incubation chamber at 37°C with 5% CO_2_ and humidity using a 30x/1.05NA silicon immersion objective lens. A 488 nm diode laser was used for excitation of Alexa Fluor 488, and a 640 nm diode laser was used for excitation of SYTO Deep Red. Laser power was adjusted between cells to maintain similar fluroesence intensity histogram between cells. All images were acquired in resonance scanning mode, with channels acquired sequentially by line on GaAsP detectors, using 6x frame averaging with a pinhole size of 2 au, a frame size of 512 x 512 pixels with pixel sizes between 0.1-0.13 µm, and z-step of between 1-1.3 µm. A gain value of 500 was used for all images. Time lapses were acquired for 5 min time intervals of 5 or 10 seconds. Prior to tracking, maximum intensity projections were generated from z-stacks. Images were denoised using the denoise.ai function in NIS Elements and then processed in FIJI by applying a median filter with a radius of 1 and running a rolling ball background subtraction with radius 30. Tracking was performed in FIJI using the TrackMate plugin with the track analysis extension to allow for automatic calculation of track length and average speed.

Spot detection was set to 1.5 µm, and tracks were inspected and corrected manually after creation. Vesicles not captured by the TrackMate plugin were tracked using FIJI’s ManualTracking plugin and then imported into TrackMate prior to running the track analysis. Distance to the cell membrane was calculated using FIJI’s outline tool with corresponding brightfield images and measurements were performed from the center of puncta to cell membrane. All data were subsequently plotted in GraphPad Prism v.9 as individual puncta. Statistical comparisons were performed using two-tailed t-tests.

### RNA pulldowns

Approximately 10 million MDA-MB-231 cells were cultured for RNA pulldown, which was conducted at 4°C. Cells were washed 3X with ice-cold PBS prior to incubation with a sufficient volume of lysis buffer 1 to cover the cell monolayer (TBS pH 7.5, 0.5% NP-40, 2.5 mM MgCl_2_, 40 u per mL RNAse inhibitor (Invitrogen), half each of complete protease and phosphatase inhibitor tablets (Roche)). After ten minutes of incubation with rocking and periodic vortexing, cell lysate was transferred to a new tube and vortexed further to complete cell lysis. Cell debris was spun down at 3,000 rpm for 5 min, and the protein content of the supernatant was assessed using a Bradford BSA standard curve. Approximately 1.5 mg protein was used per pulldown replicate. Cell lysates were pre-cleared of any non-specific streptavidin-binding proteins by incubating with 60 µL of MagneSphere paramagnetic particles (Promega, Madison, WI) for 30 min with rotation. The supernatant was removed after the beads were pulled down using a magnet stand. Beads were then washed 5X with TBS. 100 µg of biotinylated EXO-Code RNA (IDT) was added to the pre-cleared lysate and incubated for 4 h with rotation. To isolate the biotinylated RNA, 60 µL magnetic beads was added per sample and incubated for 30 min with rotation. The proteins bound to these beads were then pulled down using a magnet stand and washed 5 times with 1 mL of pulldown buffer 2 (TBS, 0.05% NP-40, RNase inhibitor, complete protease and phosphatase inhibitor tablet). After the final wash, buffer 2 was removed and beads were resuspended in 6.5 µL of ultra-clean nuclease-free water and transferred to a new tube. 1 µL of 1M DTT and 2.5 µL of 4x LDS protein loading buffer (Invitrogen) were added to the bead mixture. These proteins were then reduced at 90°C for 10 min and loaded into a 4-12% bis-tris gel (Life Technologies). This gel was run for 5 min at 150 V in 1x MOPS buffer and stained using Coomassie blue (Bio-Rad) for 1 h with rocking. Gels were then destained with methanol:acetic-acid:water in 5:1:4 ratio for 1 h at room temperature with rocking. Gels were then washed with nuclease-free water and protein bands excised for mass spectrometry.

### Mass spectrometry

After band excision, proteins were reduced and alkylated with dithiothreitol (DTT) and iodoacetamide (IAA), respectively (Sigma-Aldrich, St. Louis, MO). Gel pieces were dehydrated with acetonitrile and digested with 10 ng/µL trypsin in 50 mM ammonium bicarbonate for 30 min at room temperature, then placed at 37°C overnight. Exosomal peptides were extracted the next day by the addition of 50% acetonitrile and 0.1% trifluoroacetic acid (TFA) and were then dried in a CentriVap concentrator (Labconco, Kansas City, MO). Peptides were desalted with homemade C18 spin columns, dried again, and reconstituted in 0.1% TFA. Peptides were injected onto a 30 cm C18 column with 1.8 µm beads (Sepax, Newark, NJ,), with an Easy nanoLC-1200 HPLC (Thermo Fisher Scientific) connected to an Orbitrap Fusion Lumos mass spectrometer (Thermo Fisher Scientific). Solvent A was 0.1% formic acid in water, while solvent B was 0.1% formic acid in 80% acetonitrile. Ions were introduced to the mass spectrometer using a Nanospray Flex source operating at 2 kV (Thermo Fisher Scientific).

Peptides were eluted off the column using a multi-step gradient that began at 3% B and held for 2 min, quickly ramped to 10% B over 6 min, increased to 38% B over 95 min, then ramped to 90% B in 5 min and held for an additional 3 min to wash the column. The gradient then returned to starting conditions in 2 min and the column was re-equilibrated for 7 min, for a total run time of 120 min. The flow rate was 300 nL/min throughout the run. The Fusion Lumos was operated in data-dependent mode with a cycle time of 2 s. The full scan was conducted over a range of 375–1400 m/z, with a resolution of 120,000 at m/z of 200, an automatic gain control (AGC) target of 4e5, and a maximum injection time of 50 ms. Peptides with a charge state between 2 and 5 were selected for fragmentation. Precursor ions were fragmented by collision-induced dissociation (CID) using a collision energy of 30 and an isolation width of 1.1 m/z. MS^2^ scans were collected in the ion trap with the scan rate set to rapid, a maximum injection time of 35 ms, and an AGC setting of 1e4. Dynamic exclusion was set to 45 s to allow the mass spectrometer to fragment lower abundant peptides.

### Bioinformatics analysis

Raw data were searched with the SEQUEST search engine within the Proteome Discoverer software platform, v2.2 (Thermo Fisher Scientific) using the UniProt human database. Trypsin was selected as the enzyme, allowing up to 2 missed cleavages, with an MS^1^ mass tolerance of 10 ppm and an MS^2^ mass tolerance of 0.6 Da. Carbamidomethyl on cysteine was selected as a fixed modification. Oxidation of methionine was set as a variable modification. Percolator was used as the FDR calculator, filtering out peptides with a q-value >0.01. Label-free quantitation was performed using the Minora Feature Detector node (Thermo Fisher Scientific), with a minimum trace length of 5. The Precursor Ions Quantifier node (Thermo Fisher Scientific) was then used to calculate protein abundance ratios using only unique and razor peptides. The pairwise-based method was employed to calculate the protein ratios, which uses a protein’s median peptide ratio to determine the protein ratio. Proteins with an abundance value within the bottom 25%, or proteins that bound all sequences equally, were removed from subsequent analyses. The remaining protein abundance values were loaded into GraphPad Prism and the color mapped to the abundance value. The proteins were then divided into groups: those that solely bound the control sequence, and those that only bound the EXO-Codes. Unique proteins were then annotated for their subcellular location using Uniprot, Panther, and Reactome pathway analysis. Only proteins twice as abundant either in the control pulldown or the EXO-Codes were used for subsequent analyses. From these annotations, the proteins were grouped by broad subcellular localization and analyzed further by functionality.

### Structure predictions and functional assessment of EXO-Probes

Sequences were synthesized with varying nucleotide linker lengths and compositions (IDT). Secondary structure predictions of all sequences were performed using the RNAfold webserver with the parameters set at determining the minimum free energy of the sequence, partition function algorithm, and avoiding isolated base pairs. Entropy calculations and minimum free energy were plotted in GraphPad Prism 9. Three micromolar amounts of these sequences were then resuspended in *in vivo* mimic buffer (140 mM KCl, 1 mM MgCl_2_, 10 mM NaH_2_PO_4_ pH 7.2, 0.05% Tween-20) and incubated with 80 nm of TO1 or TO3 for 30 min at room temperature. Fluorescence readings were conducted using a Spectramax plate reader in a black-walled clear bottom plate (Nunc), with the excitation/emission for TO1 and TO3 being 510/535 nm and 637/658 nm, respectively. Data are presented as mean ± standard deviation with background fluorescence from buffer subtracted. All data were analyzed using GraphPad prism. These sequences were then electroporated into their respective cell lines and enrichment analysis conducted as described above.

### Plasmid generation of EXO-Probes

RNA Mango control and EXO-Probe gene fragments were synthesized (IDT) and cloned into the shRNA-expressing plasmid vector pSLQ1615 (Addgene, Watertown, MA). Plasmids were transformed into *E. coli* (New England Biolabs, Ipswich, MA) and isolated with the Zymopure II Midi Prep kit following the manufacturer’s protocol (Genesee Scientific, San Diego, CA). 4 µg plasmid was digested for 4 h at 37°C with *BstX*I and *Xho*I (New England Biolabs). Products were run on a 1% agarose gel for 30 min at 120 V, and digested vector was isolated using the Promega Wizard SV gel extraction and PCR cleanup kit (Promega). Products were then ligated overnight at 16°C following the NEB T4 DNA ligase protocol with 50 ng of pSLQ615. These ligations were transformed into the *E. coli* strain DH5a using NEB’s chemically-competent cell transformation protocol using ampicillin as the resistance marker. Positive colonies were grown overnight in LB medium supplement with ampicillin, and DNA was isolated using the Zyppy plasmid miniprep kit (Zymo Research). DNA was sent for Sanger sequencing (Genewiz, South Plainfield, NJ) with the following sequencing primers: FP mU6 5’-CAGCACAAAAGGAAACTCACC-3’, RP Puro 5’-GTGGGCTTGTACTCGGTCAT-3’. Clones positive for inserted fragment were used for downstream experiments.

### Transfection of EXO-Probes

100,000 MDA-MB-231 cells were seeded into one well of a 12-well plate 24-h prior to plasmid transfection. 4 µg of plasmid was transfected with 4 µL P3000 regent and 6 µL Lipofectamine 3000 as per scaled recommendations in the manufacturer’s protocol (Thermo Fisher Scientific). Medium containing lipid complexes was removed 4 h post transfection and replaced with exosome-depleted phenol-free DMEM + 10% FBS + 2 mM L-glutamine. Cells were incubated for 48 h post-transfection, and the medium collected for exosome harvest via ultracentrifugation as described above. Cells were then trypsinized and resuspended in 1 mL PBS + 0.5% BSA. Cells were analyzed on the Attune NxT Flow cytometer for relative mCherry expression (excitation/emission 587/610) under the yellow excitation laser within the 610/620 emission filter channel YL2 (Thermo Fisher Scientific). Cell populations were first gated using forward and side scatter channels and then analyzed using side scatter (y-axis) against YL2 intensity (x-axis, mCherry fluorescence).

### RNA Mango single vesicle flow cytometry

sEVs were harvested via ultracentrifugation as described above and resuspended in 20 nM-filtered PBS. These sEVs were then quantified using NTA, and protein content was determined using an equivalent Bradford standard curve. The sEVs were then incubated with a 2:1 mass ratio of TO-3 or TO-1 (ABM Technologies) for 30 min at room temperature. sEVs were then cleaned using EXO-Spin molecular weight cutoff columns (Life Technologies) and eluted in 20 nM filtered PBS. Samples were run on the Amnis ImageStream at the lowest speed possible (maximum of 600 events per second). Lasers on the ImageStream were set to the following powers unless otherwise noted: 405 nm – 150 mW, 488 nm – 100 mW, and 658 nm – 110 mW. Magnification was set to 60x, with the numerical aperture set at 0.9 and the flow core size reduced to 7 µm. TO-3 was excited using the 658 nm diode laser and analyzed on channel 11 with an emission capture range of 660-740 nm. TO-1 was excited using the 488 nm solid-state laser and data collected on channel 2 with and emission capture range between 480-560 nm. All data were collected alongside channel 6 (darkfield SSC) and two brightfield channels (1 & 9) to accurately compare fluorophores detected by both cameras of the instrument. 100 nm fluorescent polystyrene beads served as size control (Thermo Fisher Scientific). Data were analyzed using IDEAS v5.2 (Luminex, Austin, TX) and FCS Express v7.0.

### Live cell imaging of EXO-Probe movement

100,000 MDA-MB-231 cells were seeded into one well of a glass 8-well coverslip designed for inverted confocal microscopy (Ibidi Cat # 80826) 24-h prior to plasmid transfection. 4 µg of plasmid was transfected with 4 µL P3000 regent and 6 µL Lipofectamine 3000 as per scaled recommendations in the manufacturer’s protocol (Thermo Fisher Scientific). Medium containing lipid complexes was removed 4 h post transfection and replaced with exosome-depleted phenol red-free DMEM + 10% FBS + 2 mM L-glutamine (Thermo Fisher). Cells were incubated with 100 nM TO-1 in HBSS starting 30 minutes prior to imaging and for the duration of the imaging period. A 488 nm diode laser was used to excited the TO-1 fluorophore used in imaging. All images were acquired in resonance scanning mode using a 30x/1.05NA silicon immersion objective lens, 6x frame averaging with a pinhole size of 2 au, a frame size of 512 x 512 pixels with pixel sizes between 0.1-0.13 µm, a z-step of between 1-1.3 µm, and gain value of 500. Laser power was adjusted between cells to maintain similar fluroescence intensity histogram between cells. Time lapses were acquired for 5 min time intervals of 5 seconds.

Prior to tracking, maximum intensity projections were generated from z-stacks. These images were denoised using the denoise.ai function in NIS Elements and then processed in FIJI by applying a median filter with a radius of 1 and running a rolling ball background subtraction with radius 30. Tracking was performed in FIJI using the TrackMate plugin with the track analysis extension to allow for automatic calculation of track length and average speed. Spot detection was set to 1.5 µm, and tracks were inspected and corrected manually after creation. Vesicles not captured by the TrackMate plugin were tracked using manual tracking in TrackMate. Trackmate data were subsequently plotted in GraphPad Prism v.9 as individual puncta. Statistical comparisons were performed using two-tailed t-tests.

### Antibody labeling of transfected cells

#### Cy5-EXO-Code

At 4-12 hours post RNA electroporation (as described above), cells were washed 3 times before being fixed in 4% paraformaldehyde for ten minutes at room temperature. Cells were washed 3 times and permeabilized for 30 minutes in 0.1% Triton-X114 in PBS. Cells were then washed 3 times and blocked with a 2-4% BSA solution in PBS for 1 hour at RT. Primary antibodies were diluted in 2% BSA in PBS as recommended by the manufacturer (rabbit anti-human YBOX1 and EEF1A1, mouse anti-human Hnrnpm and tubulin, and goat anti-human Annexin A2), and incubated at 4 degrees C for 16 hours. Cells were then washed 5 times and incubated with secondary antibodies at a 1:2000 dilution in 2% BSA for 30 minutes at RT. Cells were washed 5 times and incubated with a working dilution of Hoechst 33342 for fifteen minutes at room temperature, prior to being washed and embedded in FluorSave mounting medium (Merck Millipore, Burlington, MA), and imaging on the Olypmus FV3000RS confocal microscope.

### Genetically-encoded EXO-Probes and TO Dye

Four hours post lipofectamine transfection (as described above), cells were washed 3 times before being fixed in 4% paraformaldehyde for ten minutes at room temperature. Cells were washed 3 times and permeabilized for 30 minutes in 0.1% Triton-X114 in PBS. Cells were then washed 3 times and blocked with a 2-4% BSA solution in PBS for 1 hour at room temperature. Primary antibodies were diluted in 2% BSA in PBS as recommended by the manufactuerer (mouse anti-human CD9 (Proteintech 60232-1-Ig), mouse anti-human CD63 (Thermofisher MEM-259), rabbit anti-human TSG101 (Abcam ab125011), and incubated at 4°C for 16 hours. Cells were then washed 5 times and incubated with secondary antibodies at a 1:2000 dilution in 2% BSA for 30 minutes at room temperature. Cells were washed 5 times and incubated with a working dilution of Hoechst 33342 for fifteen minutes at room temperature, prior to being washed and embedded in FluorSave mounting medium, and imaging on the Olypmus FV3000RS.

All images were acquired in resonance scanning mode, with channels acquired sequentially by line on GaAsP detectors, using 6x frame averaging with a pinhole size of 0.95 au, a frame size of 1024 x 1024 pixels and z-steps between 1-2.5 µm. A 405 nm diode laser was used for nuclear imaging of all samples. For the EXO-codes, a 488 nm diode laser, 561 nm diode laser, and a 640 nm diode laser were used for Hoechst, secondary mAbs (488/561) and Cy5, respectively. For the EXO-Probes, the TO dye was imaged with the 488 nm diode laser, and secondary mAbs were imaged with the 561/640 nm diode lasers. All imaging was conducted using a 60x silicon oil immersion objective lense with NA of 1.4. Co-localization analysis was done first by drawing a region of interest (ROI) around the cell of interest. The images were then split into separate channels, and analyzed via the imageJ plugin Coloc2 as described here^68^. All co-localization analysis was done with raw data files with no modification to the contrast threshold prior to calculation. Pearson’s correlation co-efficients were compiled from representative cells within no less than n=3 chamber slide wells and averaged. Data were analyzed in FIJI and GraphPad Prism, with Student’s two-tailed t-test used for statistical comparisons between groups.

### EXO-Probe secretion quantification

EXO-Probes were electroporated into MDA-MB-231 cells at a ratio of 10 µg RNA to 330,000 cells as described above. These cells were plated in triplicate into single wells of a 12-well plate in phenol-free DMEM + 10% FBS + 2 mM L-glutamine. Cells were cultured for 48 h post-electroporation, and sEVs were harvested via ultracentrifugation as described above. sEVs were diluted 1:100 in 20 nm PBS and quantified by NTA. Fixed amounts of sEVs were serially diluted in buffer (140 mM KCl, 1 mM MgCl_2_, 10 mM NaH_2_PO_4_ pH 7.2, 0.05% Tween-20) and lysed in 0.1% tween-20 for 30 min at room temperature. Samples were vortexed for three 10s pulses and incubated in a 1:1 dilution of WB buffer + 50 nM TO-3 for fifteen minutes at RT. Fluorescence values were measured using a Quibit 4 fluorometer (Thermo Fisher Scientific), and a fluorescence plate reader (Gen5). Comparisons against NTA values used the average across the three replicates for linear regression analysis.

### Statistics and reproducibility

All experiments for quantitative analysis and representative images were reproduced with similar results at minimum in triplicate unless otherwise stated. Data sets with normal distributions were analyzed with unpaired Student’s two-tailed *t*-tests to compare two groups. One-way ANOVA was used to evaluate the difference between three or more groups. Co-localization correlation analysis was performed using the Spearman’s rank correlation method. *P* < 0.05 was considered statistically significant with denominations as \*\*\*\**p* < 0.0001, \*\*\**p* < 0.001, \*\**p* < 0.01, \**p* < 0.05. All statistical analyses were performed using GraphPad Prism 9.0 software.

## Supplementary Information

**Supplementary Figure 1:**
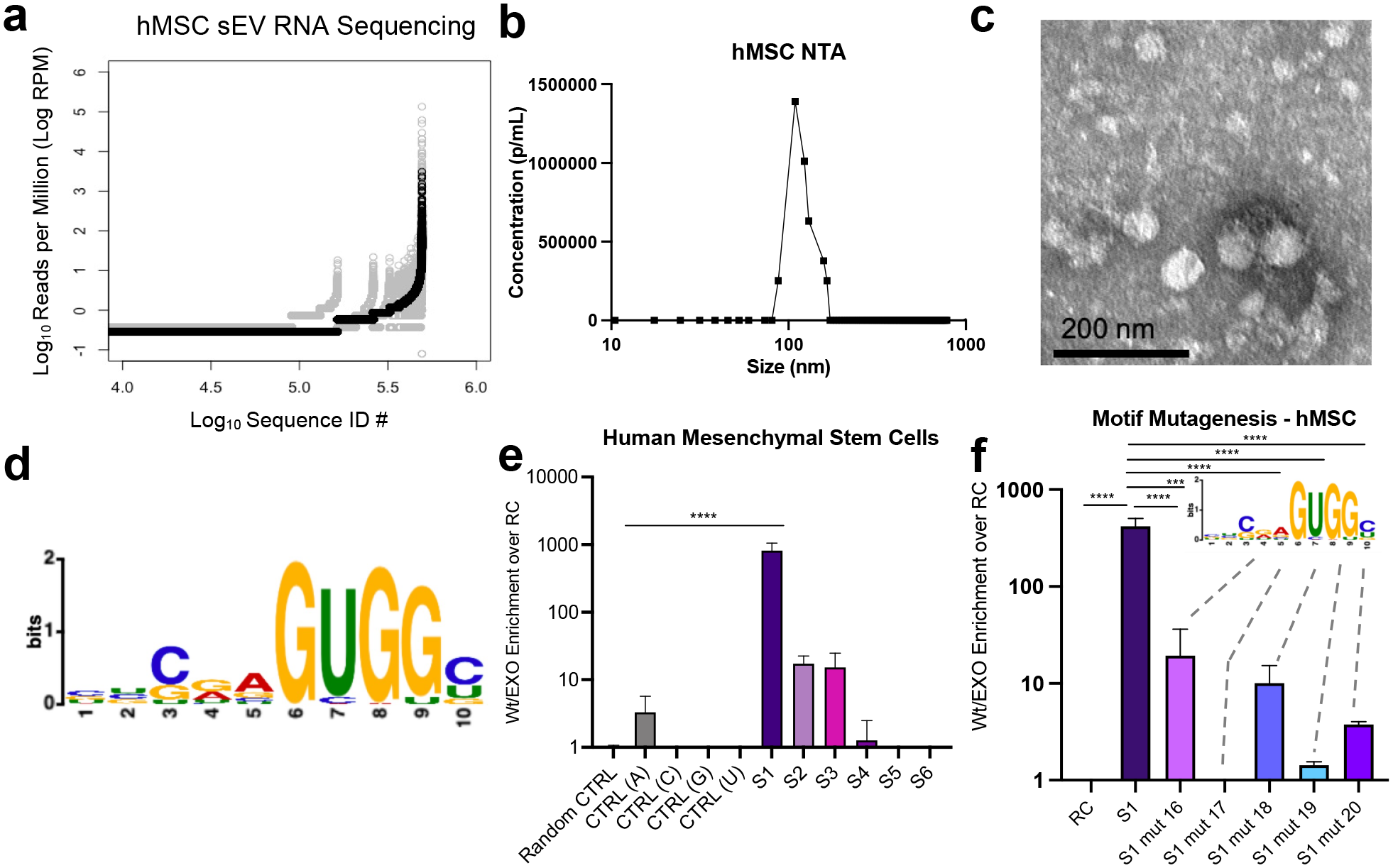
EXO-Code selection in human mesenchymal stem cells (hMSCs). **(a)** Visualization of next-generation sequencing data compiled as sequence counts versus log-normalized reads per million, with each dot representing an individual and unique sequence. Rounds 3-7 plotted in gray, and round 10 plotted in black. The overlap in sequence ID number at the upper right of the graph indicates the same unique sequences enriching into sEVs in higher amounts as the selection procedure went into later rounds. **(b)** Representative NTA plot with mean size of sEVs as 105 nm after differential ultracentrifugation. **(c)** Representative TEM image of MSC sEVs with scale bar set to 200 nm. **(d)** Bioinformatic motif derived from top 200 sequences uncovered in MSC sEVs. **(e)** Quantitative RT-PCR graph of the top six hMSC EXO-Codes as determined from panel **(a)**. **(f)** Five key motif nucleotide positions were mutated as in the M1 data. Statistical comparisons were performed using one-way ANOVA, \*\*\*\**p* < 0.0001, \*\*\**p* < 0.001, \*\**p* < 0.01, \**p* < 0.05.

**Supplementary Figure 2:**
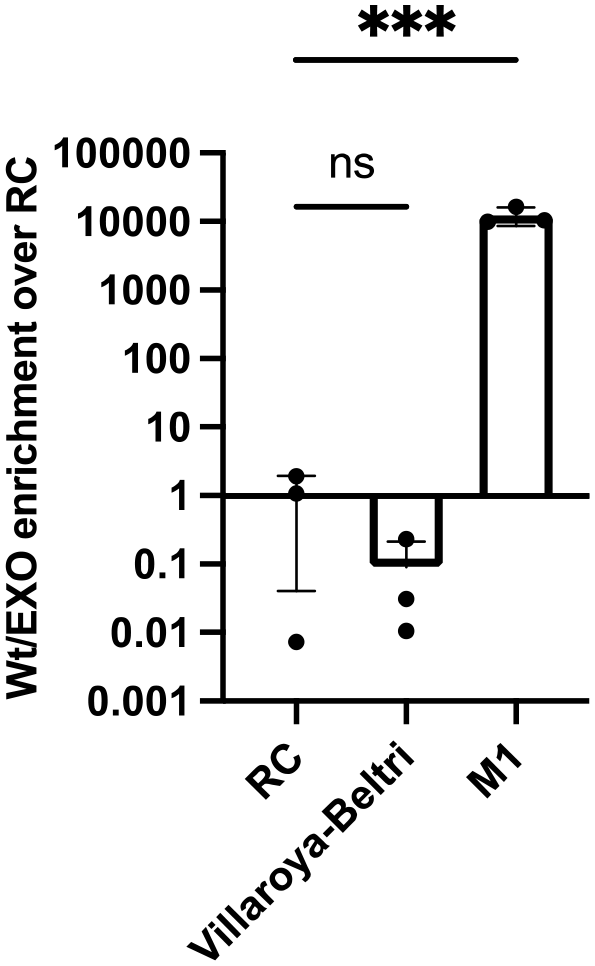
Top performing MDA EXO-Code compared against RC and 20-mer repeat of literature-reported EXO-motif. Statistical comparisons were performed using one-way ANOVA, \*\*\*\**p* < 0.0001, \*\*\**p* < 0.001, \*\**p* < 0.01, \**p* < 0.05.

**Supplementary Figure 3:**
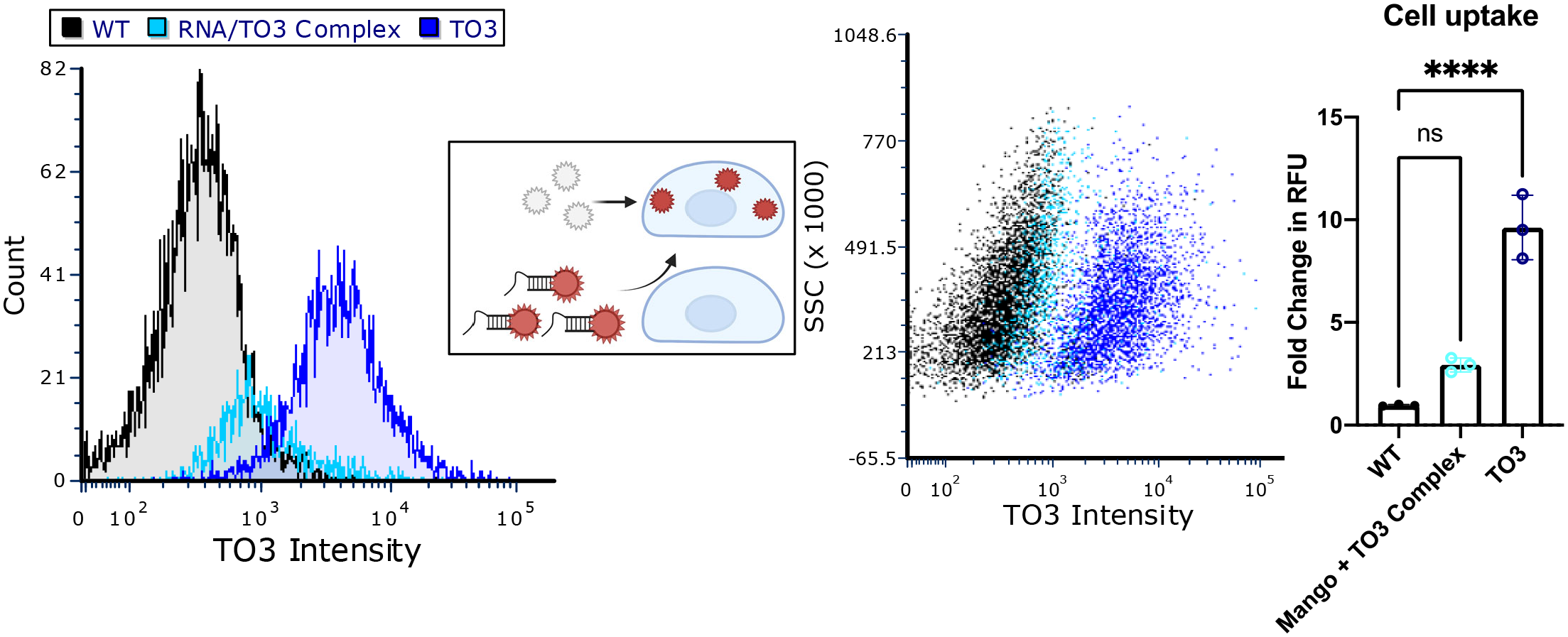
Flow cytometry plots of cellular uptake and permeability of either TO-3 alone (dark blue) or TO3 complexed with RNA (light blue). Graphical depiction of proposed mechanism. TO-3 without RNA complex is capable of permeating cells but becomes cell impermeable upon binding RNA Mango aptamer. Data were analyzed using FCS Express and GraphPad Prism. Data are presented as mean +/-standard deviation. Statistical comparisons were performed using one-way ANOVA, \*\*\*\**p* < 0.0001, \*\*\**p* < 0.001, \*\**p* < 0.01, \**p* < 0.05.

**Supplementary Figure 4:**
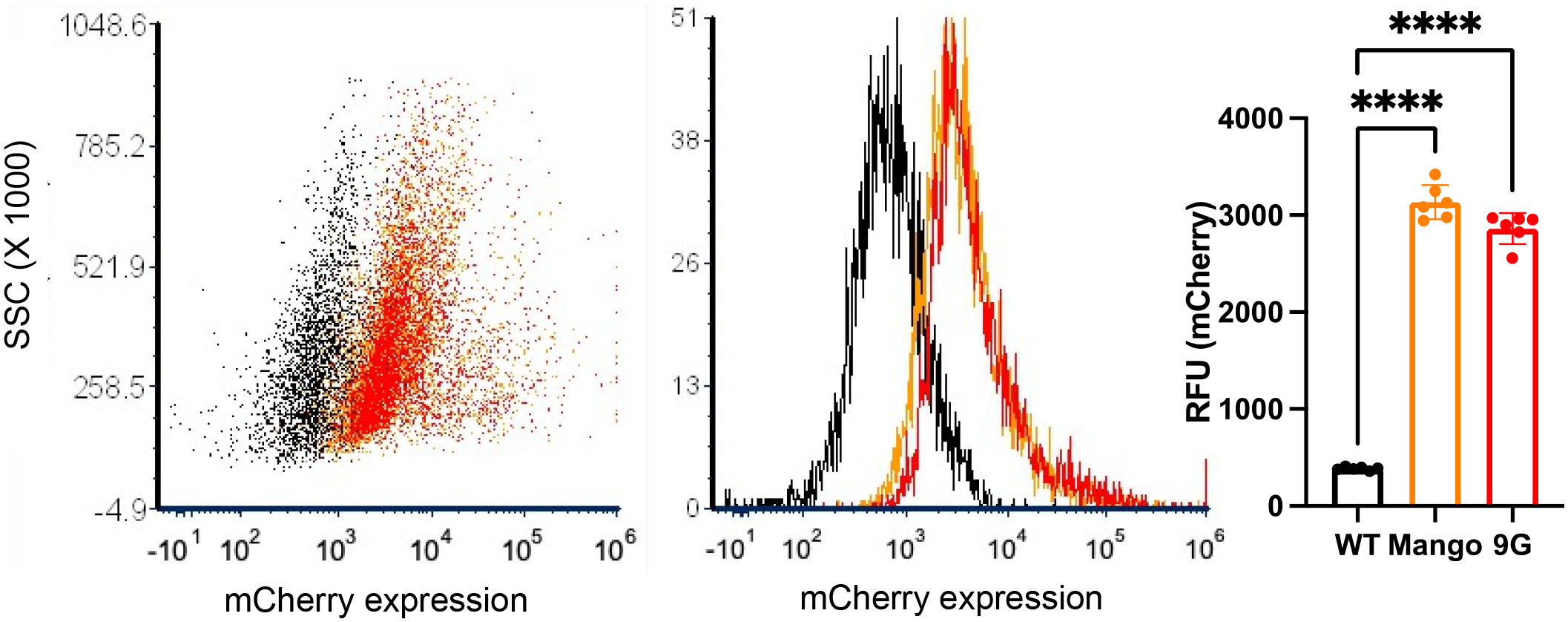
Verification of plasmid transfection with mCherry reporter. Flow cytometry plots of mCherry expression (x-axis) after transfection with pSLQ1651-EXO Probe (dark orange) and RNA mango (light orange). Data was acquired on an attune nXt (Thermo Fisher). Data were analyzed using FCS Express v.7.0 and GraphPad Prism. Data are presented as mean +/-standard deviation. Statistical comparisons were performed using one-way ANOVA, \*\*\*\**p* < 0.0001, \*\*\**p* < 0.001, \*\**p* < 0.01, \**p* < 0.05.

**Supplementary Figure 5:**
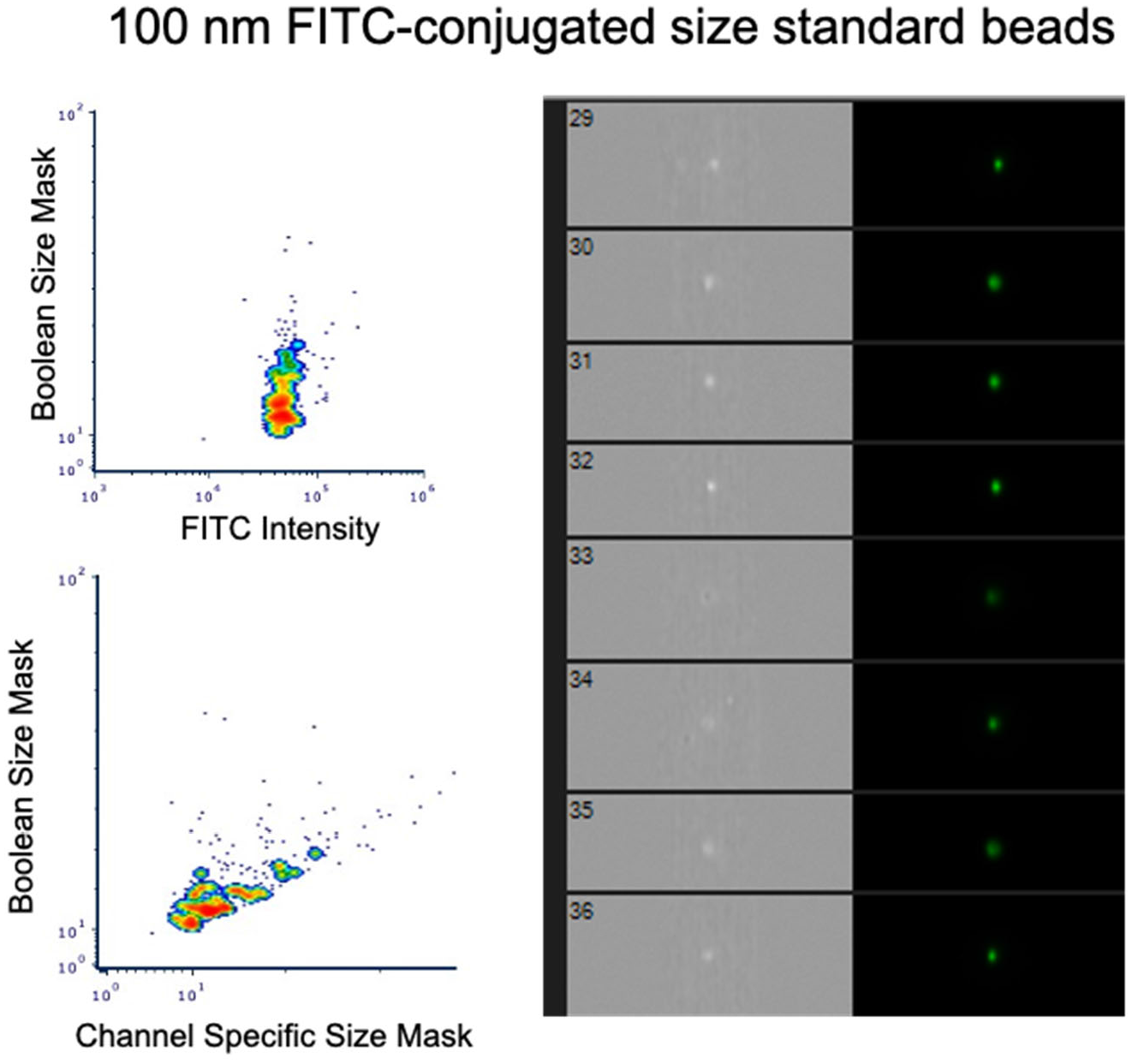
100 nM polystyrene size standard beads imaged on the Imagestream mark II hybrid flow cytometer. (Upper left) Density plot depicting uniform FITC intensity and narrow size distribution of the beads using a Boolean mask calculated from darkfield (SSC) and brightfield channels. (Bottom left) Correlation between the Boolean size mask and the fluorescent channel size mask (Channel 2 – FITC) indicating that the fluorescence does not affect size measurement. These beads were used to calibrate the machine prior to imaging and quantifying fluorescence on sEVs used in subsequent studies. Beads were resuspended in sterile-filtered dPBS and run at the slowest flow speed, 60x magnification and an 8 µM core size. Data were analyzed in both FCS Express and IDEAS (Amnis-specific flow software).

**Supplementary Table 1.**
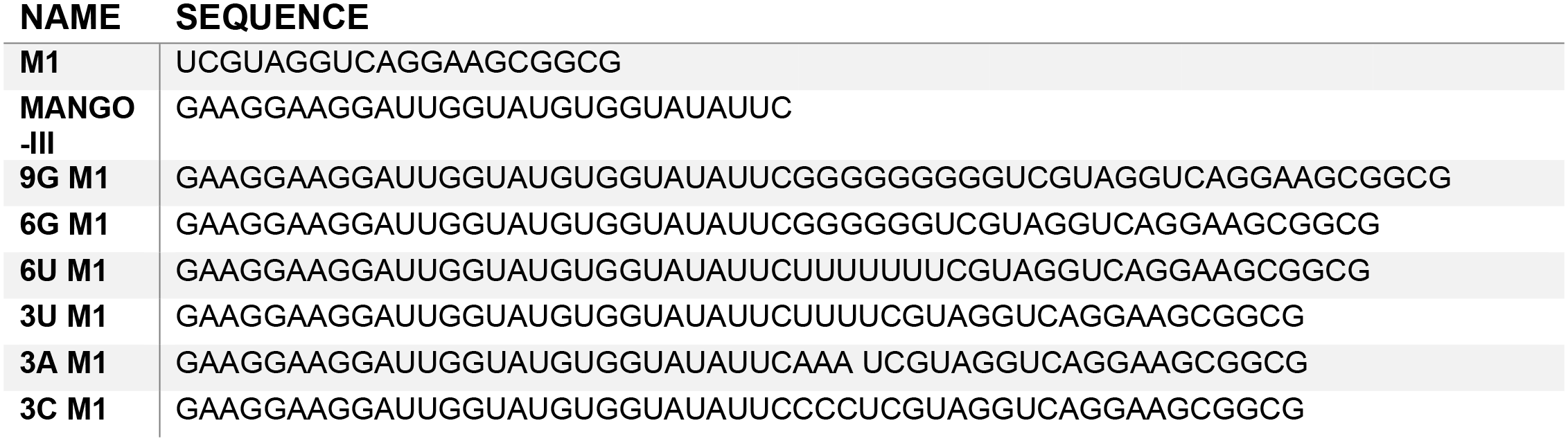
Sequences of all EXO-Probes used for structure prediction and cellular enrichment. RNA oligos were synthesized by Integrated DNA technologies (IDT) and resuspended in nuclease free water prior to use.

**Supplementary Table 2:**
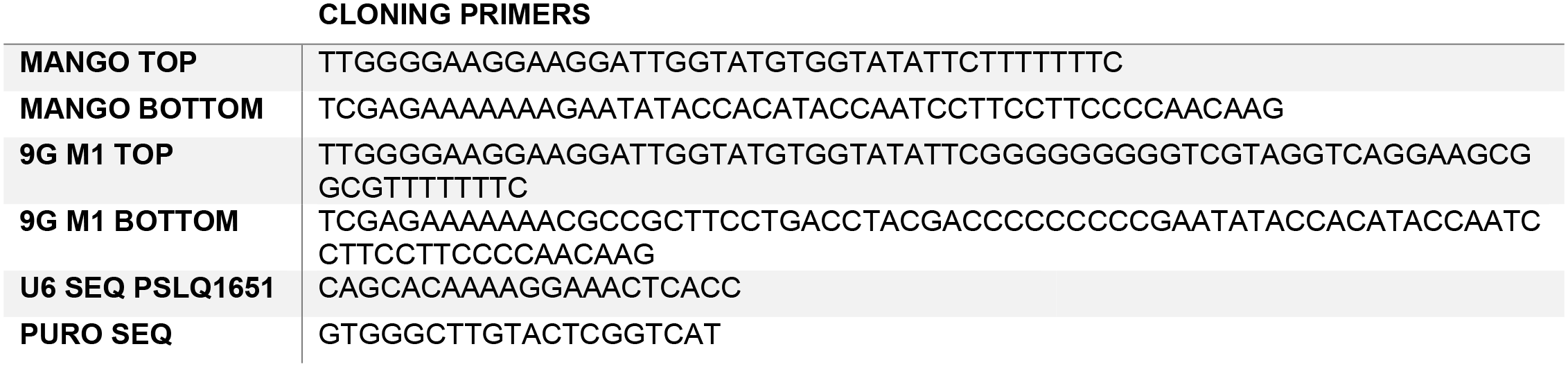
Genetically encoded EXO-Probe sequences. DNA oligonucleotides were ordered from IDT. Oligonucleotides used for cloning were annealed following the manufacturer’s protocol. Sanger sequencing was performed with the U6 and PURO sequencing primers specified above as recommended for the PSLQ615 vector.

## References

1. Théry, C. Exosomes: secreted vesicles and intercellular communications. F1000 Biol Rep 3, 15 (2011).

2. Raposo, G. & Stoorvogel, W. Extracellular vesicles: exosomes, microvesicles, and friends. J. Cell Biol. 200, 373–383 (2013).

3. Ferguson, S. W. & Nguyen, J. Exosomes as therapeutics: The implications of molecular composition and exosomal heterogeneity. J. Control. Release 228, 179–190 (2016).

4. Kishore, R. & Khan, M. More Than Tiny Sacks: Stem Cell Exosomes as Cell-Free Modality for Cardiac Repair. Circ. Res. 118, 330–343 (2016).

5. Shukla, V. C. et al. Reciprocal Signaling between Myeloid Derived Suppressor and Tumor Cells Enhances Cellular Motility and is Mediated by Structural Cues in the Microenvironment. Adv. Biosys. 4, e2000049 (2020).

6. Valadi, H. et al. Exosome-mediated transfer of mRNAs and microRNAs is a novel mechanism of genetic exchange between cells. Nat. Cell Biol. 9, 654–659 (2007).

7. Ferguson, S., Kim, S., Lee, C., Deci, M. & Nguyen, J. The Phenotypic Effects of Exosomes Secreted from Distinct Cellular Sources: a Comparative Study Based on miRNA Composition. AAPS J. 20, 67 (2018).

8. Ferguson, S. W. et al. The microRNA regulatory landscape of MSC-derived exosomes: a systems view. Sci. Rep. 8, 1419 (2018).

9. Schulz, E. et al. Biocompatible bacteria-derived vesicles show inherent antimicrobial activity. J. Control. Release 290, 46–55 (2018).

10. Sullivan, J. A. et al. Treg-Cell-Derived IL-35-Coated Extracellular Vesicles Promote Infectious Tolerance. Cell Rep. 30, 1039–1051.e5 (2020).

11. Ruan, S. et al. Extracellular vesicles as an advanced delivery biomaterial for precision cancer immunotherapy. Adv Healthc Mater e2100650 (2021). doi:10.1002/adhm.202100650

12. Jeng, S. C. Y., Chan, H. H. Y., Booy, E. P., McKenna, S. A. & Unrau, P. J. Fluorophore ligand binding and complex stabilization of the RNA Mango and RNA Spinach aptamers. RNA 22, 1884– 1892 (2016).

13. Autour, A. et al. Fluorogenic RNA Mango aptamers for imaging small non-coding RNAs in mammalian cells. Nat. Commun. 9, 656 (2018).

14. Hoshino, A. et al. Tumour exosome integrins determine organotropic metastasis. Nature 527, 329– 335 (2015).

15. Sung, B. H., Ketova, T., Hoshino, D., Zijlstra, A. & Weaver, A. M. Directional cell movement through tissues is controlled by exosome secretion. Nat. Commun. 6, 7164 (2015).

16. Temoche-Diaz, M. M. et al. Distinct mechanisms of microRNA sorting into cancer cell-derived extracellular vesicle subtypes. Elife 8, (2019).

17. Bonacquisti, E. E. & Nguyen, J. Connexin 43 (Cx43) in cancer: Implications for therapeutic approaches via gap junctions. Cancer Lett. 442, 439–444 (2019).

18. Chevillet, J. R. et al. Quantitative and stoichiometric analysis of the microRNA content of exosomes. Proc. Natl. Acad. Sci. USA 111, 14888–14893 (2014).

19. Takov, K., Yellon, D. M. & Davidson, S. M. Confounding factors in vesicle uptake studies using fluorescent lipophilic membrane dyes. J Extracell Vesicles 6, 1388731 (2017).

20. Dehghani, M., Gulvin, S. M., Flax, J. & Gaborski, T. R. Exosome labeling by lipophilic dye PKH26 results in significant increase in vesicle size. BioRxiv (2019). doi:10.1101/532028

21. Corso, G. et al. Systematic characterization of extracellular vesicle sorting domains and quantification at the single molecule - single vesicle level by fluorescence correlation spectroscopy and single particle imaging. J Extracell Vesicles 8, 1663043 (2019).

22. Lai, C. P. et al. Dynamic biodistribution of extracellular vesicles in vivo using a multimodal imaging reporter. ACS Nano 8, 483–494 (2014).

23. Hikita, T., Miyata, M., Watanabe, R. & Oneyama, C. Sensitive and rapid quantification of exosomes by fusing luciferase to exosome marker proteins. Sci. Rep. 8, 14035 (2018).

24. Verweij, F. J. et al. Live Tracking of Inter-organ Communication by Endogenous Exosomes In Vivo. Dev. Cell 48, 573–589.e4 (2019).

25. Sung, B. H. et al. A live cell reporter of exosome secretion and uptake reveals pathfinding behavior of migrating cells. Nat. Commun. 11, 2092 (2020).

26. Verweij, F. J. et al. Quantifying exosome secretion from single cells reveals a modulatory role for GPCR signaling. J. Cell Biol. 217, 1129–1142 (2018).

27. Zhao, X. et al. Exosomes as drug carriers for cancer therapy and challenges regarding exosome uptake. Biomed. Pharmacother. 128, 110237 (2020).

28. Horibe, S., Tanahashi, T., Kawauchi, S., Murakami, Y. & Rikitake, Y. Mechanism of recipient cell-dependent differences in exosome uptake. BMC Cancer 18, 47 (2018).

29. Shurtleff, M. J., Temoche-Diaz, M. M., Karfilis, K. V., Ri, S. & Schekman, R. Y-box protein 1 is required to sort microRNAs into exosomes in cells and in a cell-free reaction. Elife 5, (2016).

30. Zhang, Z., Qin, Y.-W., Brewer, G. & Jing, Q. MicroRNA degradation and turnover: regulating the regulators. Wiley Interdiscip Rev RNA 3, 593–600 (2012).

31. Zietzer, A., Werner, N. & Jansen, F. Regulatory mechanisms of microRNA sorting into extracellular vesicles. Acta Physiol (Oxf) 222, (2018).

32. Villarroya-Beltri, C. et al. Sumoylated hnRNPA2B1 controls the sorting of miRNAs into exosomes through binding to specific motifs. Nat. Commun. 4, 2980 (2013).

33. Santangelo, L. et al. The RNA-Binding Protein SYNCRIP Is a Component of the Hepatocyte Exosomal Machinery Controlling MicroRNA Sorting. Cell Rep. 17, 799–808 (2016).

34. Momen-Heravi, F. et al. Current methods for the isolation of extracellular vesicles. Biol. Chem. 394, 1253–1262 (2013).

35. Schudt, G., Kolesnikova, L., Dolnik, O., Sodeik, B. & Becker, S. Live-cell imaging of Marburg virus-infected cells uncovers actin-dependent transport of nucleocapsids over long distances. Proc. Natl. Acad. Sci. USA 110, 14402–14407 (2013).

36. Burckhardt, C. J. & Greber, U. F. Virus movements on the plasma membrane support infection and transmission between cells. PLoS Pathog. 5, e1000621 (2009).

37. Schudt, G. et al. Transport of Ebolavirus Nucleocapsids Is Dependent on Actin Polymerization: Live-Cell Imaging Analysis of Ebolavirus-Infected Cells. J. Infect. Dis. 212 Suppl 2, S160–6 (2015).

38. Bhagwat, A. R. et al. Quantitative live cell imaging reveals influenza virus manipulation of Rab11A transport through reduced dynein association. Nat. Commun. 11, 23 (2020).

39. Greber, U. F. & Way, M. A superhighway to virus infection. Cell 124, 741–754 (2006).

40. Katrukha, E. A. et al. Probing cytoskeletal modulation of passive and active intracellular dynamics using nanobody-functionalized quantum dots. Nat. Commun. 8, 14772 (2017).

41. Statello, L. et al. Identification of RNA-binding proteins in exosomes capable of interacting with different types of RNA: RBP-facilitated transport of RNAs into exosomes. PLoS One 13, e0195969 (2018).

42. Hessvik, N. P. & Llorente, A. Current knowledge on exosome biogenesis and release. Cell Mol. Life Sci. 75, 193–208 (2018).

43. Reggiori, F. & Pelham, H. R. Sorting of proteins into multivesicular bodies: ubiquitin-dependent and -independent targeting. EMBO J. 20, 5176–5186 (2001).

44. Shurtleff, M. J. et al. Broad role for YBX1 in defining the small noncoding RNA composition of exosomes. Proc. Natl. Acad. Sci. USA 114, E8987–E8995 (2017).

45. Piper, R. C. & Katzmann, D. J. Biogenesis and function of multivesicular bodies. Annu. Rev. Cell Dev. Biol. 23, 519–547 (2007).

46. Willis, G. R., Kourembanas, S. & Mitsialis, S. A. Toward Exosome-Based Therapeutics: Isolation, Heterogeneity, and Fit-for-Purpose Potency. Front. Cardiovasc. Med. 4, 63 (2017).

47. Chen, T.-M. et al. hnRNPM induces translation switch under hypoxia to promote colon cancer development. EBioMedicine 41, 299–309 (2019).

48. Lin, C.-Y., Beattie, A., Baradaran, B., Dray, E. & Duijf, P. H. G. Contradictory mRNA and protein misexpression of EEF1A1 in ductal breast carcinoma due to cell cycle regulation and cellular stress. Sci. Rep. 8, 13904 (2018).

49. Boratkó, A., Péter, M., Thalwieser, Z., Kovács, E. & Csortos, C. Elongation factor-1A1 is a novel substrate of the protein phosphatase 1-TIMAP complex. Int. J. Biochem. Cell Biol. 69, 105–113 (2015).

50. Hagiwara, K., Katsuda, T., Gailhouste, L., Kosaka, N. & Ochiya, T. Commitment of Annexin A2 in recruitment of microRNAs into extracellular vesicles. FEBS Lett. 589, 4071–4078 (2015).

51. Iparraguirre, L. et al. Circular RNA profiling reveals that circular RNAs from ANXA2 can be used as new biomarkers for multiple sclerosis. Hum. Mol. Genet. 26, 3564–3572 (2017).

52. Cawte, A. D., Unrau, P. J. & Rueda, D. S. Live cell imaging of single RNA molecules with fluorogenic Mango II arrays. Nat. Commun. 11, 1283 (2020).

53. Bartys, N., Kierzek, R. & Lisowiec-Wachnicka, J. The regulation properties of RNA secondary structure in alternative splicing. Biochim. Biophys. Acta Gene Regul. Mech. 1862, 194401 (2019).

54. Sabarinathan, R. et al. The RNAsnp web server: predicting SNP effects on local RNA secondary structure. Nucleic Acids Res. 41, W475–9 (2013).

55. Buratti, E. & Baralle, F. E. Influence of RNA secondary structure on the pre-mRNA splicing process. Mol. Cell. Biol. 24, 10505–10514 (2004).

56. Vohhodina, J. et al. BRCA1 binds TERRA RNA and suppresses R-Loop-based telomeric DNA damage. Nat. Commun. 12, 3542 (2021).

57. Crescitelli, R. et al. Distinct RNA profiles in subpopulations of extracellular vesicles: apoptotic bodies, microvesicles and exosomes. J Extracell Vesicles 2, (2013).

58. Morales-Kastresana, A. et al. Labeling extracellular vesicles for nanoscale flow cytometry. Sci. Rep. 7, 1878 (2017).

59. Willms, E. et al. Cells release subpopulations of exosomes with distinct molecular and biological properties. Sci. Rep. 6, 22519 (2016).

60. Dunn, K. W., Kamocka, M. M. & McDonald, J. H. A practical guide to evaluating colocalization in biological microscopy. Am. J. Physiol. Cell Physiol. 300, C723–42 (2011).

61. de Jong, O. G. et al. A CRISPR-Cas9-based reporter system for single-cell detection of extracellular vesicle-mediated functional transfer of RNA. Nat. Commun. 11, 1113 (2020).

62. Dominska, M. & Dykxhoorn, D. M. Breaking down the barriers: siRNA delivery and endosome escape. J. Cell Sci. 123, 1183–1189 (2010).

63. Patel, S. et al. Naturally-occurring cholesterol analogues in lipid nanoparticles induce polymorphic shape and enhance intracellular delivery of mRNA. Nat. Commun. 11, 983 (2020).

64. Cardarelli, F. et al. The intracellular trafficking mechanism of Lipofectamine-based transfection reagents and its implication for gene delivery. Sci. Rep. 6, 25879 (2016).

65. Maruyama, T., Nara, K., Yoshikawa, H. & Suzuki, N. Txk, a member of the non-receptor tyrosine kinase of the Tec family, forms a complex with poly(ADP-ribose) polymerase 1 and elongation factor 1alpha and regulates interferon-gamma gene transcription in Th1 cells. Clin. Exp. Immunol. 147, 164–175 (2007).

66. Shelke, G. V., Lässer, C., Gho, Y. S. & Lötvall, J. Importance of exosome depletion protocols to eliminate functional and RNA-containing extracellular vesicles from fetal bovine serum. J Extracell Vesicles 3, (2014).

67. FASTAptamer Users Guide. At <https://usermanual.wiki/Document/FASTAptamerUsersGuide.1565872219/view>

68. Coloc 2. at <https://imagej.net/plugins/coloc-2>

